# Sequence-dependent co-condensation of Lsr2 with DNA elucidates the mechanism of genome compaction in *Mycobacterium tuberculosis*

**DOI:** 10.1101/2025.01.06.631453

**Authors:** Thejas Satheesh, Rohit Kumar Singh, Prakshi Gaur, Sneha Shahu, Saminathan Ramakrishnan, Mansi Srivastava, Shreyasi Neogi, Sandeep Choubey, Mahipal Ganji

## Abstract

The xenogeneic silencer protein Lsr2 from *Mycobacterium tuberculosis* plays a critical role in its survival and pathogenesis. Lsr2 is a nucleoid-associated protein that interacts with DNA *in vivo* and regulates many genes. Purified Lsr2 forms nucleoprotein filaments with DNA molecules leading to highly compacted DNA conformations. However, the physical mechanism underlying Lsr2-mediated DNA compaction, resulting in gene regulation, remains elusive. We employed a combination of biochemical assay, single-molecule imaging, and molecular dynamics simulations to investigate the governing principles of Lsr2-mediated DNA compaction. We show that, while Lsr2 alone undergoes phase separation, addition of DNA substantially lowers the required concentration for its phase separation. Strikingly, our single-molecule and simulation data establish that Lsr2 forms condensates with long stretches of AT-rich DNA, providing strong evidence for sequence-dependent co-condensation. This observation is contrary to the classical view of sequence-dependent binding of individual protein molecules to DNA, our findings rather suggest that protein-DNA co-condensates ‘sense’ the average binding energy landscape. We present a physical model for Lsr2-mediated DNA compaction and gene regulation, describing a novel mechanism for NAP-mediated genome organization in bacteria.

## Introduction

The specific organization of bacterial genome and the tight regulation of genes are crucial for controlling cell fate dynamics for survival and infection^1^. The Nucleoid-Associated Proteins (NAPs) are a diverse and abundant group that are expressed in growth phase-dependent manner.^2,3^ In addition to their role in maintaining nucleoid architecture, NAPs are responsible for modulating the accessibility of DNA to transcriptional machinery thereby directly regulating global gene expression^4,5^. However, the physical mechanism by which NAPs compact the genome and regulate gene expression in bacteria remains elusive.

In this report, we sought to probe the mechanism of action of a crucial NAP, called Lsr2 from *Mycobacterium tuberculosis* (*Mtb*). Lsr2 assumes a key role in regulating the expression of genes linked to virulence^6^. For instance, Lsr2 regulates the expression of iniBAC operon which is known for imparting multi-drug tolerance in *Mtb* upon the front-line antibiotic drugs treatment. Studies have shown that *Mtb* lacking Lsr2 were unable to adapt to extreme oxygen levels, resulting in compromised long-term survival^7^. Additionally, Lsr2 was shown to be essential for disease pathology and the establishment of chronic infections in the mouse models of tuberculosis^7^. Thus, Lsr2 dictates the ability of *Mtb* to adapt to the adverse conditions encountered during infection and helps in its persistence within the host. Hence, understanding the molecular mechanisms of Lsr2-DNA interactions will pave the way for controlling bacterial pathogenicity^8^.

Our current understanding of Lsr2-DNA interactions largely stems from biochemical, structural, and force-spectroscopy studies^9–12^. Structurally, Lsr2 possesses a C-terminal DNA-binding domain and N-terminal dimerization and oligomerization domain (Fig. 1a and 1b)^12^. Chromatin-Immunoprecipitation and sequencing (ChIP-seq) analysis showed that Lsr2 exhibits greater preference for AT-rich sequences, encompassing hundreds of genes^10^. In agreement with these studies, biochemical studies indicated that Lsr2 exhibits concentration-dependent DNA binding, forming multiple protein-DNA complexes, with a preference for AT-rich sequences^9^. Furthermore, single-molecule magnetic tweezer experiments demonstrated that Lsr2 binding to DNA generates a rigid Lsr2 nucleoprotein complex, leading to compaction of DNA^11^.

**Figure 1:**
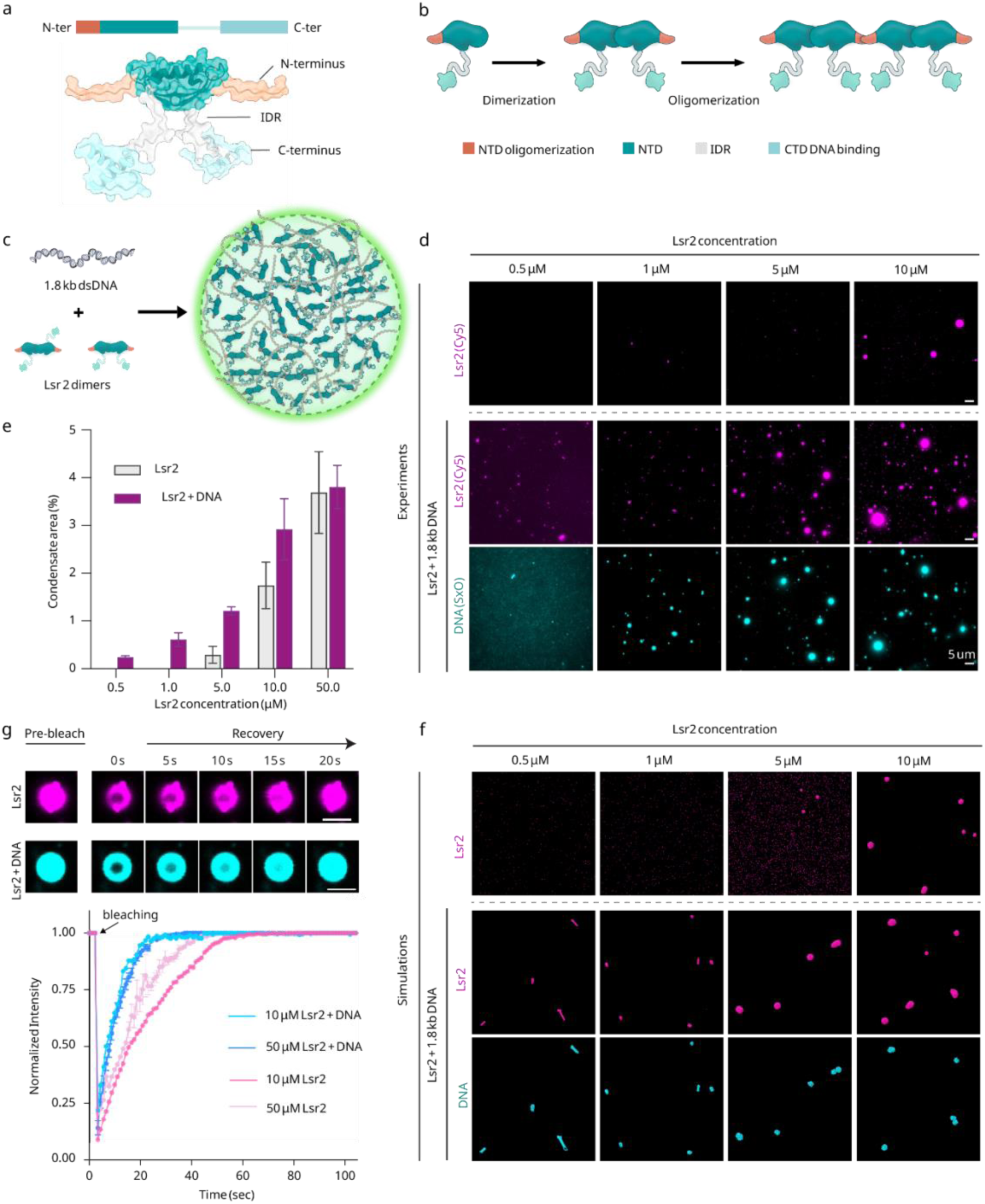
Lsr2 and DNA undergo liquid-liquid phase-separation together. a) AlphaFold prediction of the full-length Lsr2 dimer. The N-terminal domain (NTD) is responsible for the oligomerization of Lsr2 dimers, and the C-terminal domain (CTD) plays a role in DNA binding. The NTD and CTD are connected by an intrinsically disordered region (IDR) spanning 20 amino acids. b) Cartoon representation of Lsr2 monomer, dimer, and oligomer. c) Schematic Lsr2-DNA co-condensation. d) Fluorescence images of Lsr2 condensates at different concentrations of Lsr2 alone (top row) and with 1.5 nM of 1.8 kb-long dsDNA (bottom row) (scale bar – 3 µm). e) Quantification of condensate area for the data in figure 1d. f) Using coarse-grained Brownian dynamics simulations (See SI), we show that our experimental findings are consistent with a simple model of protein condensation in the presence of DNA, driven by protein-protein and protein-DNA interactions. g) (top) Fluorescence images showing the FRAP of Lsr2 (above) and Lsr2-DNA condensates. (bottom) Plot showing the recovery of normalized fluorescence intensity after photobleaching.

Recently, single-cell imaging of *M. smegmatis* revealed the role of Lsr2 in chromosome organization and cell cycle^13^. The real-time analysis of fluorescently tagged Lsr2 showed the formation of one or two prominent foci, accompanied by various minor complexes. As the cell cycle progresses, the Lsr2 focus splits into two and move to the opposite poles. Since DNA replication happens during the same time, Lsr2 nucleoprotein complexes were speculated to undergo dynamic assembly and disassembly facilitating the organization of newly replicated DNA. While these *in vitro* and *in vivo* studies revealed a variety of interactions that Lsr2 displays with DNA, the physical principles governing the formation of Lsr2-DNA complexes, resulting in genome organization and large-scale gene regulation, are not well-understood.

In this manuscript, we aimed to dissect the physical mechanism of Lsr2-mediated DNA compaction. To this end, we implemented a range of biochemical, biophysical, and single-molecule methods in combination with coarse-grained molecular dynamics simulations to delineate the physical mechanism behind Lsr2-mediated DNA compaction and its potential role as a xenogeneic regulator. We observed that purified Lsr2 protein alone forms phase-separated condensates. Interestingly, the addition of DNA reduced the concentration required condensates formation. Single-molecule visualization under a Total Internal Reflection Fluorescence (TIRF) microscope revealed that Lsr2 and DNA co-condense in a sequence-dependent manner, with Lsr2 preferentially localizing on and compacting stretches of AT-rich DNA regions. In contrast to the long-standing view that individual proteins binding to their DNA sequence motifs, we find that Lsr2 proteins bind collectively to form co-condensates along extended DNA segments characterized by high-density clusters of motifs with moderately high or high AT-content. Such DNA sequence-dependent co-condensation of Lsr2 and DNA can sequester many genes, thereby revealing a potential gene silencing mechanism.

## Results

### Lsr2 and DNA together undergo phase separation

To shed light on the physical mechanism of genome organization by Lsr2, we perform biochemical characterization of the purified protein. We observed that purified Lsr2 forms a turbid solution at 4 °C. The solution turned clear when it was further diluted, transferred into a high salt buffer. Upon microscopy inspection of the turbid solution, we observed the presence of phase separated condensates (Fig. 1c). Next, we probed the concentration-dependence of these Lsr2 condensates by fluorescently labeling and subsequent visualization under fluorescence microscope. Our observations revealed that Lsr2 phase-separates at a concentration of ∼10 µM and the area occupied by the condensates increased with Lsr2 concentration (Fig. 1d and 1e)^14^. The condensate size increased as a function of Lsr2 concentration (Fig. 1e). Such Lsr2 condensates underwent fusion and showed Fluorescence Recovery After Photobleaching (FRAP) to the original fluorescence levels, the hallmarks of phase separated condensates (Fig. 1g and Supplementary Figure 1a). Interestingly, when we added DNA to the solution, Lsr2 formed condensates at a much lower concentration of about 1 µM (Fig. 1d). Moreover, DNA and Lsr2 co-localized in these condensates. Once again Lsr2 displayed FRAP to the original fluorescence levels, albeit at a higher rate than Lsr2 alone (Fig. 1g and Supplementary Figure 1c, d). Disrupting the ionic interactions by increasing the salt concentration caused the condensates to dissolve, demonstrating the reversibility of condensate formation (Supplementary Figure 1b).

We performed both Electrophoretic Mobility Shift Assay (EMSA) and Atomic Force Microscopy (AFM) to visualize the Lsr2-DNA complexes. These observations clearly demonstrated the formation of higher-order Lsr2-DNA complexes (Supplementary figure 2 a-c).

While our observations are consistent with the structural properties of Lsr2 (Fig. 2a), which are often suggested as prerequisites for a biomolecule to phase separate, i.e., i) the presence of a low complexity sequence, which results in intrinsically disordered region (IDR), connecting its dimerization (N-terminus) and DNA-binding domains (C-terminus)^10^, and ii) Lsr2’s ability to oligomerize^12^, we intend to probe how each of these aspects contribute towards the formation of Lsr2 condensates.

**Figure 2:**
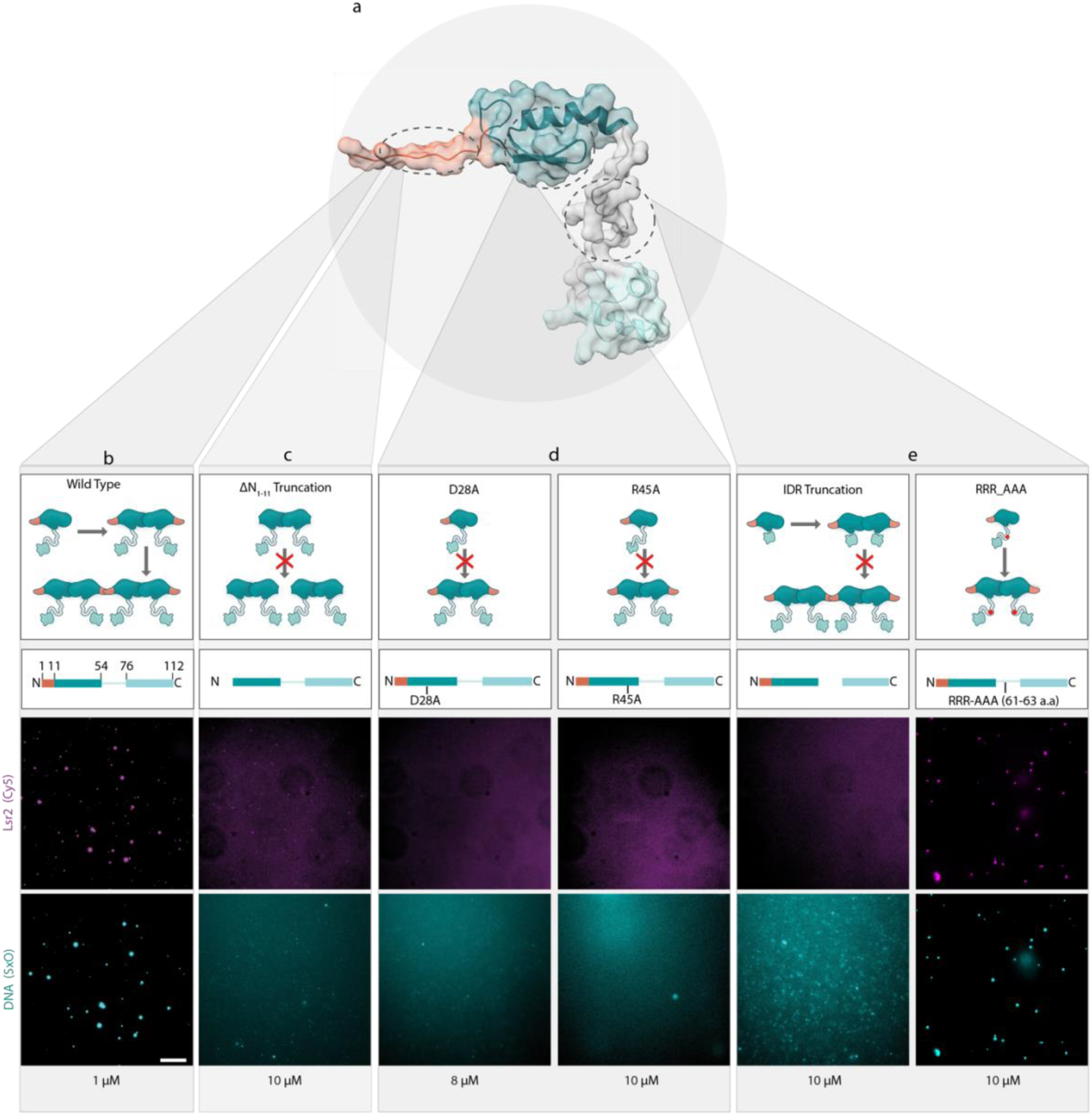
Oligomerization domain and intrinsically disordered region are necessary for phase-separation. a) Alfafold2^15^ predicted structure of WT Lsr2 monomer. b) Cartoon representation of formation of Lsr2 dimer and oligomers from monomer. The corresponding fluorescence images of Lsr2-Cy5 and DNA(SxO) of a solution consisting of both Lsr2 and DNA. c) Cartoon showing N-terminal eleven amino acid truncate of Lsr2 dimer which cannot form oligomers. The corresponding fluorescence images of Lsr2- Cy5 and DNA(SxO) of a solution consisting of both Lsr2 and DNA. d) Individual point mutations leading to abolition of dimerization. The corresponding fluorescence images of Lsr2-Cy5 and DNA(SxO) of a solution consisting of both Lsr2 and DNA. e) Truncation of IDR and mutation of RRR to AAA within IDR and its corresponding fluorescence images of Lsr2 and DNA. Scale bar in all the microscopy images is 10 µm

### Both multivalent binding and intrinsically disordered regions are required for phase-separation of Lsr2

We probed how multivalent interactions of Lsr2 resulting from its oligomerization dictates its condensation. Notably, Lsr2 exists as a dimer which can bind DNA at two sites and its ability to oligomerize via its N-terminal interactions enables multivalent DNA binding. Additionally, Lsr2 carries a long IDR enriched with arginine and glycine amino acids which are implicated in weak protein-protein interactions^16,17^.

We systematically dissected the role of each of these characteristics of Lsr2 in its ability to phase-separate both in the presence and absence of DNA. First, we tested the role of dimerization (Fig. 2d). A previous study had shown that D28A or R45A mutations completely abolish dimerization of Lsr2^9^. In our experiments, both the dimerization mutants failed to form condensates, indicating that dimerization is key to phase separation of Lsr2 (Fig. 2d). Without forming dimers, the N-terminal oligomerization domain of the monomers can only interact with each other, thereby limiting the oligomeric state to divalent interactions. To further unravel the role of multivalent interactions, we generated another mutant by truncating eleven amino acids from the N-terminus (ΔN_1-11_), which is thought to be responsible for oligomerization^12^. The ΔN_1-11_ mutant also failed to undergo condensation (Fig. 2c). Taken together, these findings indicate that the ability to oligomerize through multivalent binding is essential for phase separation.

Next, we examined the role of IDR that links the N- and C-terminus of Lsr2 on its condensation. To this end, we created two Lsr2 mutants. In the first one, consecutive arginine amino acids (from 61-63) of the IDR were mutated to alanine (RRR mutant). We observed that the RRR mutant required a much higher Lsr2 concentration to phase separate in the presence of DNA as compared to the WT Lsr2 (Fig. 2e). The fraction of condensed area was comparable between 1 µM of WT Lsr2 + DNA (0.7 % ± 0.2%) and 10 µM of RRR Lsr2 + DNA (0.6% ± 0.2%) (Supplementary Figure 1e).

To explore this further, we generated another mutant in which a major part of the IDR (amino acids 58-73) was truncated, creating a Lsr2_ΔIDR_ version. In accordance with our hypothesis, the Lsr2_ΔIDR_ failed to undergo phase separation (Fig. 2e). These observations lead us to conclude that both multivalent binding via oligomerization, and weak molecular interactions mediated via the IDR region are necessary for the phase separation of Lsr2.

A recent live-cell imaging study of *M. smegmatis* revealed that Lsr2 appears as a punctate *in vivo*^13^. It remains unclear how Lsr2 can function as a global gene regulator while existing as a single punctate^7,10^. In view of our observation that Lsr2 can undergo phase-separation, we hypothesized that Lsr2 and DNA together can form phase separated condensates, which could explain its mechanism of action in bacteria. Specifically, Lsr2 forms condensates with preferred regions of DNA by looping it inside the condensate, a phenomenon known as sequence-dependent protein-DNA co-condensation^18^, which can elegantly explain the regulation of specific set of genes. To test this hypothesis, we turned to the direct visualization of Lsr2 condensation on DNA using single-molecule imaging experiments.

### Two-color single-molecule imaging reveals co-condensation of Lsr2 and DNA

We employed a single-molecule assay^19^ to visualize the formation of Lsr2 condensates on DNA in real-time (Fig. 3a). Individual double-stranded λ-phage DNA molecules of 48.5-kilo basepairs (kb) labelled with biotin on both ends were immobilized on a PEG-passivated glass surface. We introduced DNA with a low buffer flow rate such that the end-to-end length of the tethered DNA is shorter than its contour length. We visualized DNA molecules under TIRF microscope after staining with an intercalating dye^19^. Upon introduction of Lsr2 into the flow cell, we observed a rapid increase in the fluorescence intensity at specific locations along the DNA, which otherwise appeared homogenous. The intensity of the fluorescent spots increased gradually at specific positions along the DNA for both double and single tethered DNA (Fig. 3b). To test our hypothesis of Lsr2-DNA co-condensation, we performed two-color simultaneous imaging of DNA and Lsr2. As expected, we observed the formation of Lsr2-DNA co-condensates along the DNA in which the fluorescence intensity of Lsr2 increased until reaching a steady state (Fig. 3c-3e). Interestingly, in addition to local compaction at specific locations, we observed coating of Lsr2 along the length of DNA which resulted in concomitant decrease in fluorescence signal from DNA (Fig. 3c-3e). The loss of DNA intensity is due to inaccessibility to the SxO dye molecules by Lsr2 coated on DNA. Taken together, these observations indicate an initial coating of Lsr2 molecules on the DNA forming nucleoprotein-filament, followed by DNA bridging and cluster formation by Lsr2 at specific locations on DNA (Fig. 3c). Notably, such condensation of Lsr2 and DNA led to a loss in DNA fluctuations which we quantified by analyzing the DNA envelope width (Fig. 3f)^20^. We observed that the envelope width is independent of end-to-end length of DNA when Lsr2 is present, indicating that Lsr2 compacts double-tethered DNA against a specific force.

**Figure 3:**
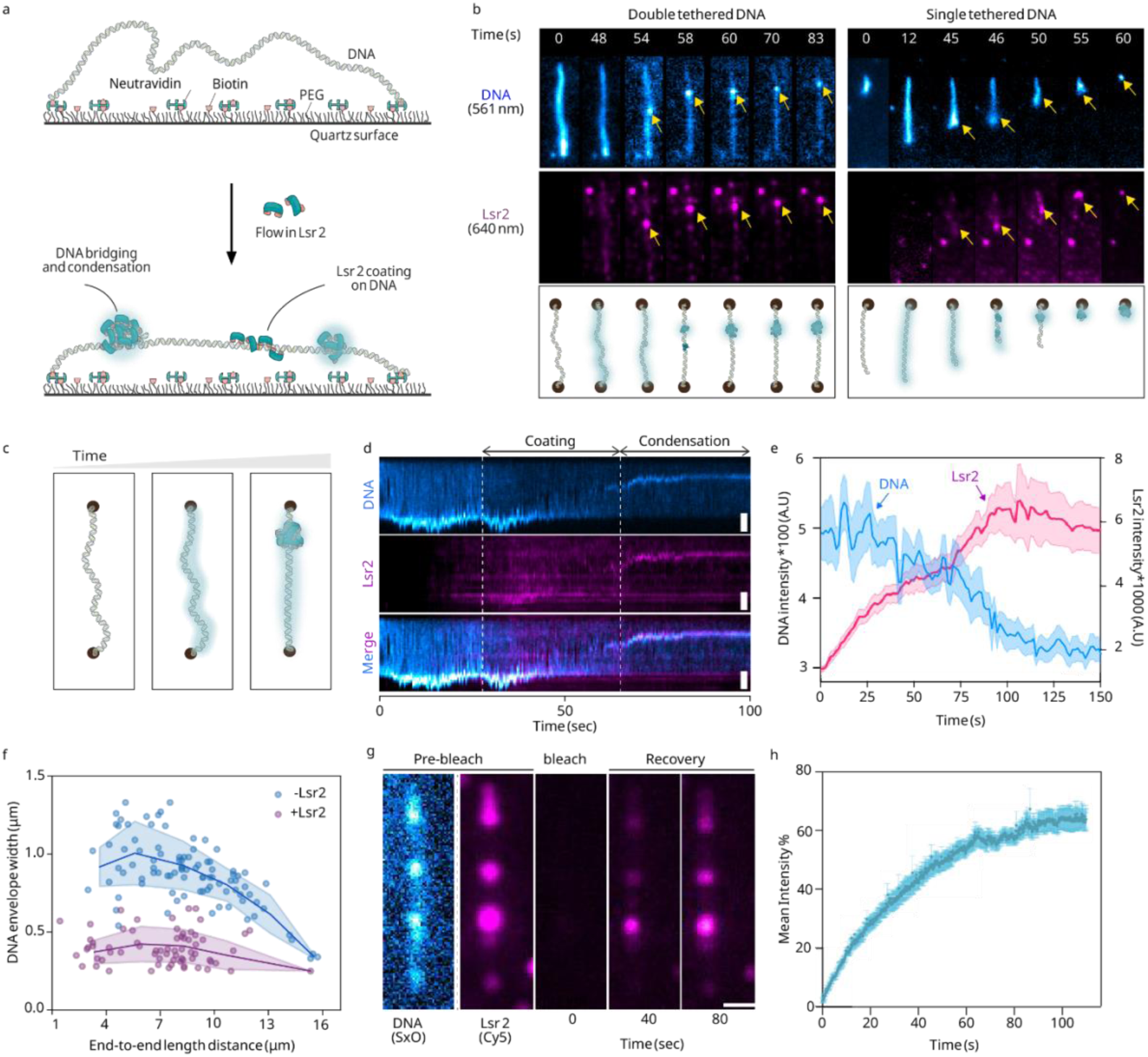
Single-molecule studies on Lsr2-DNA interactions. a) Side view representation of the single molecule assay. The DNA molecule is tethered onto the PEG passivated surface by biotin- streptavidin linkages. Lsr2 is then flowed in, and the interactions are monitored in real time. b) A series of snapshots illustrates both the DNA channel (depicted in blue) and the Lsr2 channel (depicted in magenta) for a double tethered DNA molecule on the left and a single tethered DNA molecule on the right. c) Schematic illustration of the sequence of events observed in the experiments that Lsr2 coating on DNA precedes compaction. d) Representative kymographs illustrate the intensity of DNA and Lsr2 over time. A decrease in DNA intensity is observed concurrently with an increase in Lsr2 intensity along the length of the DNA molecule. e) Intensity plots of Lsr2 and DNA across the entire DNA molecule confirm the coating of Lsr2 along the DNA. f) DNA envelope width as a function of the end-to-end distance of DNA measured in the absence (red) and presence of Lsr2 (blue). g) FRAP of Lsr2-DNA condensates. h) Quantification of FRAP in figure (3g).

To further investigate the fluidity of the DNA-Lsr2 co-condensates formed on tethered DNA molecules, we once again performed FRAP (Fig. 3g). We first photobleached the fluorescence of Lsr2 molecules using high laser power under TIRF illumination which we then replaced with labeled Lsr2. We observed recovery of fluorescence at the exact same positions where the clusters were seen before photobleaching, albeit slower than in the condensates shown in figure 1g (Fig. 3h). The slow recovery is likely due to the lower concentration of proteins used in the single- molecule experiments than in the biochemical analysis. Additionally, we observed multiple clusters merging and coalescing over time that appeared along the length of DNA (Supplementary Figure 3b). These collective observations further support the emergence of Lsr2-DNA co- condensates.

Additionally, we performed similar single-molecule experiments using the Lsr2 mutant variants (Supplementary Fig. 3c–h). The observations from these experiments align with the findings of the biochemical analysis, as shown in figure 2.

The number of Lsr2-DNA condensates formed on the tethered DNA decreased with the end-to-end distance of the DNA and ceased to show any compaction closer to contour length of DNA (Supplementary Figure. 4a). This data indicates that the Lsr2-DNA co-condensation is sensitive to the force applied across the length of DNA, at forces closer to 1 pN corresponding to the contour length of DNA, Lsr2 ceased to co-condense with DNA. While the condensates tended form towards at specific positions along the λ-DNA (Supplementary Figure. 4b, c), suggesting that DNA sequence might play a role on condensate formation. Owing to the fact that Lsr2 preferentially binds AT-rich regions of DNA^10^, we speculated that the condensates’ positions were dictated by the AT-content of the DNA.

### Lsr2 forms sequence-dependent co-condensates with DNA at AT-rich regions

To dissect the sequence dependency of Lsr2-DNA co-condensation, we prepared two DNA constructs: i) 18 kb DNA with relatively homogeneous AT-content (Fig. 4a), and ii) 21 kb long DNA with a 3 kb long AT-rich segment in the middle of construct-i (Fig. 4e) and labeled them with Cy5 at one of the ends (Fig. 4a, e). Imaging Cy5 provided us with the directionality of the tethered DNA, which enabled us to explore the role of DNA sequence on the position of Lsr2-DNA co-condensates. We observed that the majority of the Lsr2 condensates (86.4 %) formed at the center of the double-tethered molecules where AT–rich sequence is present (Fig. 4f and 4g, Supplementary Figure. 4d-iv). In sharp contrast, in the control construct lacking AT-rich sequence, Lsr2 doesn’t show any preference for condensate formation (Fig. 4b and 4c, Supplementary Figure. 4d-iv). In another control, when we replaced the high AT-content of the 21 kb-long DNA construct with moderate AT content, we once again observed a homogeneous pattern of condensate formation along the DNA (Supplementary Figure. 4g). Our analysis of the intensity profiles of individual DNA molecules revealed a clear correlation (Pearson’s correlation of 0.9) between the DNA sequence and Lsr2-mediated DNA compaction (Supplementary figure 3a). To rule out the possibility of DNA-tethering on the position of condensate formation, we prepared another Cy5-labeled DNA construct with the AT-rich segment placed on the other side of the Cy-labeled end. On this construct, we observed condensates forming at the AT-rich segment again, confirming that AT-content dictates the position of Lsr2 condensates (Supplementary Figure 4f). In summary, Lsr2 specifically condenses AT-rich segments of DNA.

**Figure 4:**
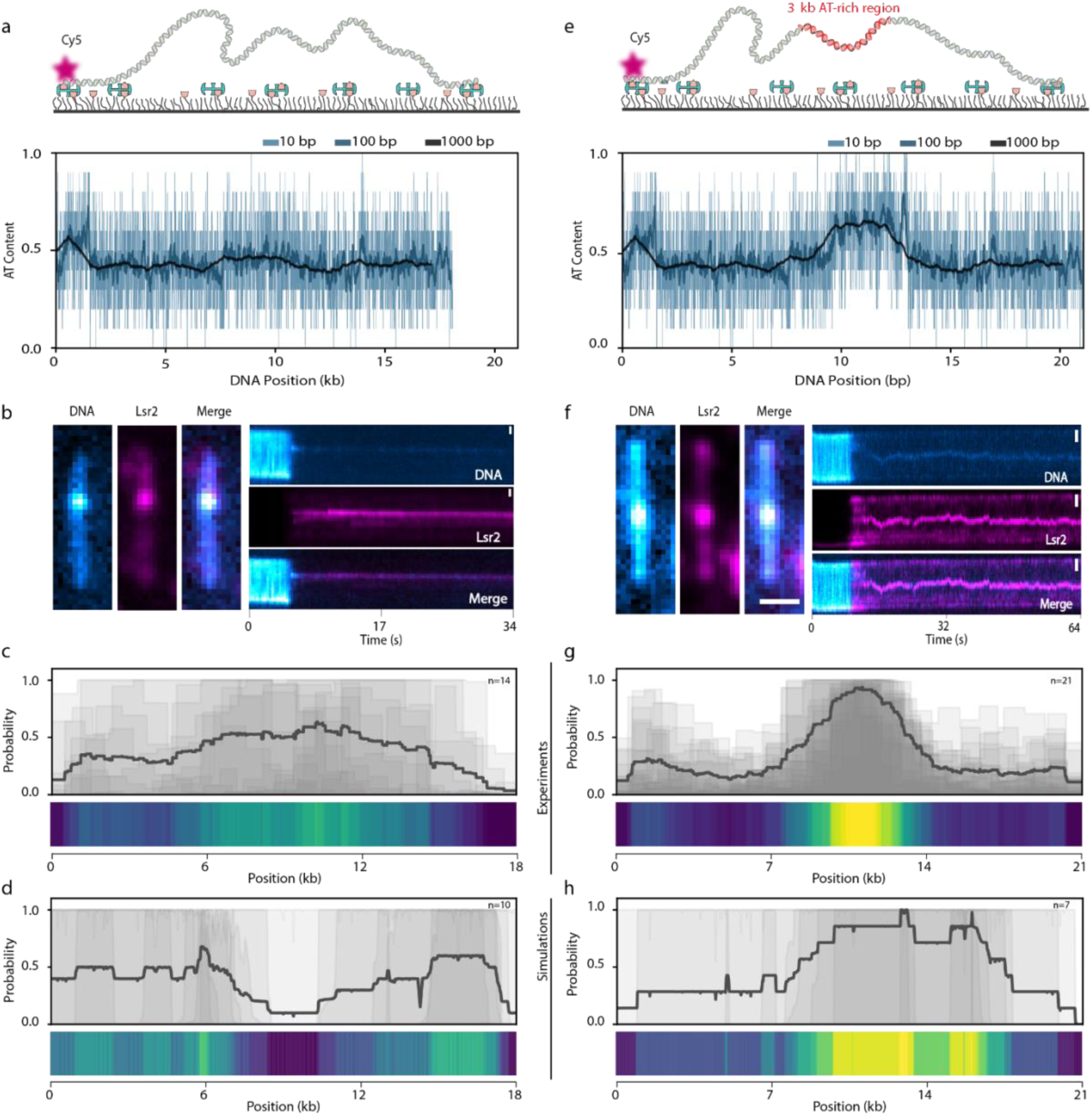
Lsr2 co-condenses on AT-rich region of DNA. a) (Top) Side-view schematic of the Cy5-labelled DNA construct used in the assay without the presence of an AT-rich region. (Bottom) The average AT-content for the DNA constructs using coarse-graining window of 10, 100, and 1000 base pairs. b) Snapshots captured post DNA compaction, depicting the DNA channel and Lsr2 channel for figure 4a. The corresponding kymographs depicting the time evolution of co- condensation are shown on the right. c) (Top) Experiment – the probability of different DNA segments observed inside the condensate is depicted along the length of DNA. (bottom) Heatmap of the probability profile. d) (Top) Simulation – the probability of different DNA segments observed inside the condensate is depicted along the length of DNA in figure 4a. (bottom) Heatmap of the probability profile. e) (Top) Side-view schematic of the DNA construct with an AT-rich segment. (Bottom) The average AT-content for the DNA constructs using coarse-graining window of 10, 100, and 1000 base pairs. f) Snapshots captured post DNA compaction, depicting the DNA channel and Lsr2 channel for DNA shown in figure 4e. The corresponding kymographs depicting the time evolution of co-condensation are shown on the right. g) (Top) Experiment – the probability of different DNA segments observed inside the condensate is depicted along the length of DNA shown in figure 4e. (bottom) Heatmap of the probability profile. d) (Top) Simulation – the probability of different DNA segments observed inside the condensate is depicted along the length of DNA corresponding to figure 4e. (bottom) Heatmap of the probability profile.

The data so far indicated that Lsr2 forms co-condensates along with DNA in a sequence-dependent manner. Notably, while individual Lsr2 molecules exhibit preferential binding to 10 bp AT-rich DNA recognition motifs^10^, the co-condensation phenomenon observed in our experiments arises from the collective behavior of many Lsr2 molecules. We sought to unravel how individual Lsr2 binding at the length scale of a few base pairs leads to the emergence of the sequence-dependent co-condensation of Lsr2 at larger kilo base pair length scales, as observed in our experiments. To this end, we developed a simple physical model of protein-DNA co-condensation that explicitly incorporates DNA sequence heterogeneity for protein binding^18^. We constructed simple physical models to shed light on our experimental findings for both i) the biochemical assay and ii) the single-molecule setup. In both these models, we represented Lsr2 proteins by spherical beads with specific binding affinities to each other, and DNA as a semiflexible polymer in a cuboidal box (Supplementary Information). Since Lsr2 footprint on the DNA is about 10 base pairs^6^, we assumed that each monomer in the polymer represents ten base pairs of DNA, and the spherical beads have identical dimensions as the monomers. While the polymer is freely suspended for the biochemical assay, we tethered the two ends of the polymer to capture the single-molecule experimental setup. In the case of the biochemical assay, we considered a homogeneous polymer with all the monomers having identical affinity for the proteins. However, to unravel the sequence-dependency of co-condensation corresponding to the single-molecule setup, we introduced a heterogeneous polymer consisting of monomers with varying affinities determined by their AT- content (Supplementary Information).

By employing Brownian dynamics simulations, we systematically explored these two models (Supplementary Information). In agreement with the observations from our biochemical assay, we found that Lsr2 alone can phase separate at about ∼10 µM concentration, while the addition of DNA led to condensation well below the saturation concentration for bulk phase separation of Lsr2 (Fig. 1f). These results suggest that protein-protein and protein-DNA interactions suffice in explaining the experimental observations. Next, we probed the sequence-dependency of co-condensation by employing our model corresponding to the single-molecule assay. Guided by the single-molecule assay, we chose polymers with the same sequence and length as in Figure 4a (Supplementary Information). Our model predicted that for the DNA polymer with relatively homogeneous AT-content, Lsr2 doesn’t show any preferred location for condensates formation (Fig. 4d). In contrast, for the 21 kb DNA with 3 kb AT-rich segment in the middle, condensates tended to form at the AT-rich sequence in the middle of the DNA in excellent agreement with the experimental observations (Pearson’s correlation coefficient of ∼0.7) (Fig. 4h). Our model allows us to connect the base pair level binding of Lsr2 to DNA with the emerging co-condensation patterns that involve many protein molecules and long stretches of DNA. A key finding of our theory-experiment dialogue is that Lsr2 proteins bind collectively to form co-condensates along extended DNA segments characterized by high-density clusters of motifs with moderately high or high AT content. This finding stands in sharp contrast to the classical view of DNA sequence-dependent binding of individual proteins for performing their function.

## Discussion

Taken together, our study provides the first experimental evidence for *DNA sequence-dependent co-condensation* of protein and DNA. We note that these observations are distinct from earlier experimental observations of sequence-dependent surface condensation^21^, sequence-independent co-condensation^22–24^, bridging-induced phase separation^25,26^ or co-condensation of proteins and nucleic acids^27–31^. Building upon these findings, we propose a mechanistic model to explain the role of Lsr2 in genome organization and as a global regulator of gene expression wherein Lsr2 undergoes sequence-dependent co-condensation with DNA. In this model, Lsr2 can bind different genomic regions in a sequence-dependent manner, pulling these regions into the co-condensate, thereby exerting control over gene expression while appearing as a punctae in the cell (Fig. 5).

**Figure 5:**
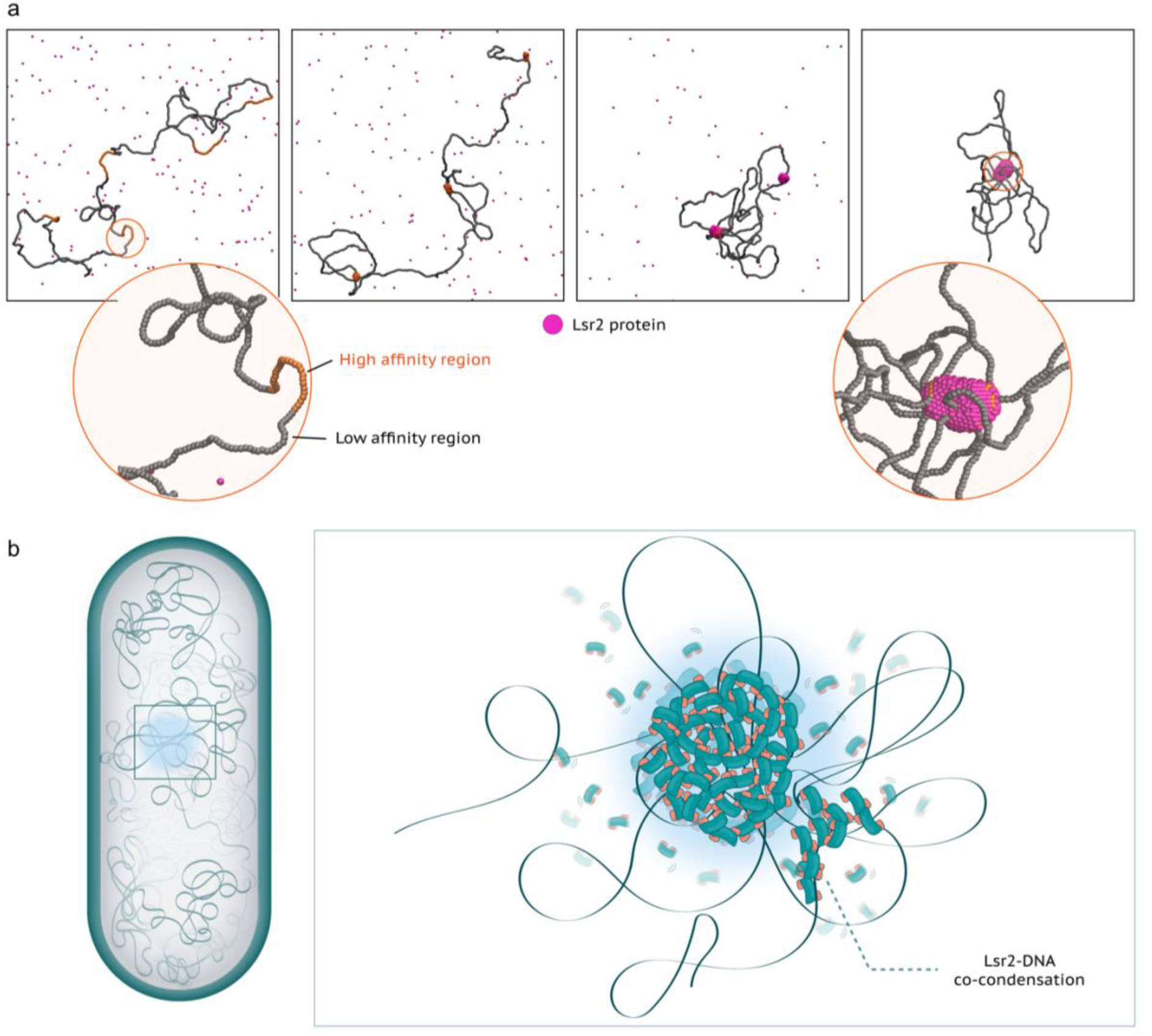
Model for Lsr2-mediated genome compaction and large-scale gene regulation. a) Simulation results of a block co-polymer of 1000 monomers (10 kb DNA). The Lsr2 proteins have an affinity of 4 k_B_T between themselves. The low affinity block of the polymer (shown in grey) interacts with Lsr2 with affinity of 0.1 k_B_T while high affinity block (shown in orange) interacts with Lsr2 with affinity of 5 k_B_T. b) According to our model, the prevailing scenario is that Lsr2 forms co-condensates by sequestering specific regions of the DNA thereby controlling the expression of a large number of genes.

While NAPs display a wide range of DNA interactions including bending, bridging or wrapping, their actual mechanism of action in folding the chromosome and regulating gene expression mechanism are unclear. An abundant protein in the stationary phase of *E. coli*, called Dps condenses DNA in a sequence-independent but rather in a topology-dependent manner^32,33^. While the chromosome conformation capture data in Mtb is lacking, an Lsr2 analog protein called Rok in *Bacillus subtilis* has been shown to mediate long-range chromosomal loops^34^. Like Lsr2, Rok is also a xenogeneic silencer protein; hence, both proteins might function via similar physical mechanisms. Interestingly, H-NS, another functional ortholog of Lsr2, is only shown to bridge DNA^35^, leading to local compaction, but so far there is no evidence for the formation of long-range loops. Nevertheless, our single-molecule imaging coupled with theoretical modeling allowed us to identify sequence-dependent co-condensation of a NAP and DNA as a novel mechanism for gene silencing in bacteria.

Bacteria carries a set of NAPs which play a variety of roles in chromatin transactions including DNA compaction and gene regulation. A comprehensive understanding of the physical mechanism behind their functioning remains elusive. Previous studies showed that Lsr2 is a functional ortholog of H-NS protein which can bridge different segments of DNA^36^. Our study revealed a physical mechanism of Lsr2-mediated DNA compaction; Lsr2 forms co-condensates with DNA. We performed an extensive biochemical analysis of a range of Lsr2 mutants to delineate the different factors dictating Lsr2-DNA co-condensation. Our study revealed that Lsr2-mediated DNA compaction results from a combined effect of multivalent binding of Lsr2 on DNA and the weak inter-molecular interactions conferred by its IDR region.

Lsr2 is a xenogeneic silencing factor from Mtb that plays a crucial role in the virulence of the bacterium. Previous Chip-Seq analyses have shown that Lsr2 controls hundreds of genes in *Mtb^10^*. Interestingly, live-cell imaging of model bacteria *M. smegmatis* showed that Lsr2 appears as clusters in bacteria^13^. Using a set of single-molecule experiments and a simple physical model, we establish that Lsr2 co-condenses with DNA specifically on AT-rich DNA sequences which are often horizontally acquired genetic elements^37^. Sequence-dependent co-condensation of a protein with DNA can allow it to simultaneously regulate a large number of genes. Indeed, our simulations confirm that high-affinity DNA regions can be sequestered into the protein-DNA co-condensate by looping out low affinity regions, offering a novel mechanism for gene regulation by NAPs in bacteria. We speculate that multicomponent phase separation involving protein and DNA, such as the one we observed, may have implications for the pathogenicity of Mtb^38^.

Our study uncovers a unique mode of DNA sequence recognition, distinct from the binding of individual protein molecules to DNA. While single-molecule binding typically occurs at the scale of a few base pairs, the formation of Lsr2-DNA co-condensates occurs over much longer stretches of DNA, spanning thousands of base pairs. This observation was confirmed experimentally and further supported by our simulations. In particular, Lsr2-DNA co-condensates preferentially form in regions of the DNA where the average binding energy is higher compared to surrounding areas. These regions may contain both weak and strong binding motifs, but their collective average affinity is stronger than in other regions of the DNA. This suggests that Lsr2 co-condensation ‘senses’ the overall binding energy landscape, rather than specifically recognizing individual motifs^10^. Our findings are particularly relevant in the context of recent experimental findings suggesting that weak affinity flanking sequences around the core DNA binding motifs of transcription factors play a crucial role in achieving their sequence specificity^39,40^. This behavior may have functional consequences, where DNA regions are selectively sequestered into the condensates based on their average sequence properties, thereby exerting control over gene expression (Figure 5).

## Acknowledgements

We acknowledge Pinaki Swain for technical discussions for executing simulations. We thank Cees Dekker for providing us DNA plasmids for single-molecule imaging experiments. We thank V. Nagaraja for kind gift of Lsr2 expression plasmid. We thank K. Muniyappa for allowing us to use their AFM facility. We acknowledge Department of Science and Technology, Ministry of Science and Technology, India DST-FIST Program funded Central Facility, Department of Biochemistry, IISc. Support from DBT/Wellcome India Alliance intermediate fellowship (IA/I/21/2/505928) to M. G. is greatly appreciated. We acknowledge funding from Department of Biotechnology (BT/PR40186/BTIS/137/3/2020). We greatly acknowledge the support from Max-Planck Institute of Immunology and Epigenetics Freiburg, Germany in terms of partner group to Mahipal Ganji lab. S.C. acknowledges the support provided by the DBT Ramalingaswami Fellowship.

## Material and Methods

### Lsr2 expression and purification

The lsr2 encoding gene was cloned into the pET-28a expression vector. This clone was kindly provided by Prof. V. Nagaraja’s Lab, MCB, IISc. An additional cysteine was introduced at 3^rd^ position from the N-terminus of the Lsr2 (Lsr2-Cys) via site-directed mutagenesis. The cysteine was inserted at third position because this residue is exposed on protein’s surface for site specific labelling with fluorophores. Briefly, the Lsr2-Cys mutant ORF was amplified from the wild-type Lsr2 clone with the forward primer Lsr2_cys_NcoI_FP containing NcoI restriction site and a codon for cysteine amino acid and reverse primer Lsr2_cys_XhoI_RP for XhoI restriction site (Table 3). The amplicon was purified after gel electrophoresis and the product was digested with NcoI (NEB R31935) and XhoI (NEB R0146L) restriction enzymes. The resulting product was then ligated with similarly digested pET-28a cloning vector using T4 DNA Ligase (NEB M0202S). The incorporation of the lsr2 gene was confirmed by sequencing. The mutants D28A, R45A, RRR_AAA (61-63), N-terminus truncation (1-52) of Lsr2-Cys were prepared using Q5 site-directed mutagenesis kit (NEB E0552S). All the clones were confirmed by sequencing. The primer sets used for these mutants are listed in Table 2.

The resulting plasmid was introduced into *E. coli* Rosetta (DE3) cells. The expressed Lsr2 gene carried a 6×His-tag fused at its N-terminus. Rosetta (DE3) cells containing the recombinant plasmid were grown at 37 °C in Lauria Broth (LB, HIMEDIA M575) media containing 50 μg/ml kanamycin (GBiosciences AB1025). After reaching the late log phase O.D. to A_600_ ∼0.5, the cells were induced with 0.3 mM of IPTG (isopropyl β-d-1 thiogalactopyranoside) (GBiosciences RC1113A) for 4 hours at 37 °C. Subsequently, the cells were harvested by centrifugation at 6000 rpm at 16 °C for 20 minutes. The resulting cell pellet was then resuspended in a lysis buffer containing 10 mM Tris-HCl pH 7.5 (Tris-Base, Sigma 77861; HCl, Fisher Scientific 29507), 1 M NaCl (SRL 3205), 1 mM DTT (DL-Dithiothreitol, SRL 17315), and 5% glycerol (Sigma G5516- 1L). The cell suspension was lysed by sonication. The cell lysate was then cleared by ultracentrifugation at 18300×g for 40 minutes at room temperature.

The cleared supernatant was applied to the Ni-NTA beads (GBiosciences 786-940) and allowed to incubate for 1 hour. The beads were washed with wash buffer 1 (10 mM Tris-HCl pH 7.5, 1.5 M NaCl, 20 mM Imidazole (SRL 61510), 5% glycerol) followed by wash buffer 2 (10 mM Tris-HCl pH 7.5, 100 mM NaCl, 50 mM Imidazole, 5% glycerol), and wash buffer 3 (10 mM Tris-HCl pH 7.5, 100 mM NaCl, 100 mM Imidazole, 5% glycerol). The protein was eluted with 500 mM Imidazole containing 10 mM Tris-HCl pH 7.5, 100 mM NaCl, and 5% glycerol. The eluted sample was applied to the Heparin beads (GBiosciences 786-842), and then the Lsr2 protein was eluted with 10 mM Tris-HCl pH 7.5, 600 mM NaCl and 5% glycerol. The protein was aliquoted stored at -80 °C.

### Fluorescent labelling of Lsr2 for imaging experiments

Lsr2 with single cysteine (Lsr2-Cys) was purified through Ni-NTA affinity chromatography followed by a similar protocol used for WT Lsr2 purification.

Purified Lsr2-Cys protein was mixed with Cy3-maleimide/Cy5-maleimide (Cytiva GEPA23031/ Cytiva GEPA15131) in a 1:3 molar ratio. The reaction mixture was incubated in the dark for 4 hours at room temperature. The excess free fluorophore was removed by buffer exchange using the Amicon ultra centrifugal filter 10 kDa MWCO (Amicon UFC5010). The filter was prepared by passing the 400 µl milli-Q and then equilibrated with the 400 µl labelling buffer (10 mM Tris-HCl, 600 mM NaCl, and 5% glycerol) by spinning at 13,000 rpm. The 100 µl protein labelling mixture was loaded onto the column with 400 µl labelling buffer, spun, and buffer exchanged to remove the free fluorophore until the flow-through became colourless. The labelling efficiency of the protein was measured approximately 20-25% by measuring the concentration ratio at 280 nm (*ɛ*_280nm_ = 13,980 M^−1^ cm^−1^) for Lsr2 and 560 nm (*ɛ*_560nm_ = 150,000 M^−1^ cm^−1^) for Cy3 or 640 nm (*ɛ*_640nm_ = 250,000 M^−1^ cm^−1^) for Cy5. We stored the protein -80°C after aliquoting.

### Atomic Force Microscopy (AFM)

AFM imaging was performed on an Agilent 5500 atomic force microscope under tapping mode. All the topographs were recorded at 1024×1024 pixel-Hz scan rate using cantilevers (NuNano Scout 70) with spring constant of 2 N/m and resonance frequency of 70 kHz. AFM topographs were processed by using Gwyddion software^41^. The AFM sample was prepared by mixing the 1.8 kb of DNA with varying concentrations of Lsr2 and incubated in a binding buffer containing 10 mM Tris-HCl pH 7.5 and 100 mM NaCl for 30 minutes at 27 °C. The control DNA sample was made by keeping all the conditions same without the Lsr2 protein. The reaction mixture was applied on the freshly peeled mica and incubated for 2.5 minutes with the adsorption buffer composed of 20 mM Tris-HCl pH 7.5, 20 mM NaCl,100 μM spermidine (SRL 17030) and 25 mM MgCl_2_ (Magnesium chloride, EMPLURA 105833) in 1:10 ratio and further rinsed with approximately 12 ml of HPLC grade water (SRL 92605), excess water removed by drying under nitrogen gas. Then, the sample was imaged under the microscope.

### Electrophoretic Mobility Shift Assay (EMSA) of Lsr2-DNA complexes

Cy5-labelled 30-mer nucleotide, linear 1.8 kb, 4.8 kb, and 6.6 kb double-stranded DNA fragments were used for the mobility shift assays. Lsr2 protein in different concentrations was mixed with double-stranded DNA and incubated for 30 minutes at 27 °C in the DNA binding buffer (10 mM Tris-HCl pH 7.5, 100 mM NaCl). The total volume of the reaction mixture was 20 µl. Following incubation, 5 µl of gel-loading dye (NEB B70248) was added, and the samples were loaded onto the 1% agarose gel (Lonza Seakem@LE Agarose 50004) in 1×Sodium Borate buffer (45 g boric acid SRL 80266; 8 g NaOH SRL 68151). The gel was run at 150 V for Cy5-labelled 30-mer DNA for 1 hour and 3 hours in the case of 1.8 kb, 4.8 kb, and 6.6 kb lengths of DNA. Once electrophoresis was done, the gels were either imaged directly in the AZURE biosystem phosphor- imager (Cy5-labelled 30-mer) or stained with EtBr (E7637) and imaged in the Gel-doc (1.8 kb, 4.8 kb, and 6.6 kb DNA).

### Preparation of biotinylated lambda DNA construct

Lambda DNA is 48.5 kb linear, double stranded DNA, native of *E. coli* lambda bacteriophage. It carries single-stranded cohesive ends of12 bp. Biotin moieties were incorporated into the cohesive ends of the Lambda phage DNA (SRL 6729) using Klenow fragment (NEB M0210L) mediated end filling in the presence of dCTP, dGTP, dATP (GBiosciences 786-460), and biotin-dUTP (Jena Bioscience NU-803-BIOX-S) at 25 °C for 15 minutes, then heat-inactivated the polymerase by incubating at 75 °C for 20 minutes. The biotin-labelled DNA was purified using a PCR clean-up kit (Monarch T1030S) to remove the excess nucleotides and polymerase. This preparation provides us with double-tether DNA molecules with a single biotin at each end. We also got single tether molecules, where biotin is present only at one end, and some molecules without biotin, which do not bind to the PEG passivated streptavidin-coated surface. The biotinylated DNA was digested using a KasI (NEB R0544S) or ApaI (NEB R0114S) restriction enzyme to generate single biotin fragments of 45.68 kb and 2.82 kb, 38.41 kb and 10.090 kb, respectively. This preparation resulted in DNA molecules with a single biotin moiety at one end.

### Preparation of DNA construct with 3 kb high AT-content in the middle

A 21 kb-long DNA was prepared by inserting a 3 kb-long 66% AT-rich region in middle of the 18 kb vector. The 3 kb-long AT-rich DNA region was amplified from Lambda phage DNA and incorporated XbaI site using forward primer 3 kb 66% AT+XbaI_FP and PciI site through reverse primer 3 kb 66% AT+PciI_RP by PCR. Amplified fragment was digested with XbaI (NEB R0145S) and PciI (NEB R0655S) restriction enzymes and ligated to the already digested 18 kb plasmid with the same enzymes. The obtained ligated plasmid was transformed into DH5-alpha competent cells to amplify it. The obtained colonies were screened for 3 kb AT-rich insert by colony PCR. ). To prepare biotinylated DNA molecules, the 21 kb plasmid with 3 kb AT-rich in the center was digested with NotI (NEB R3189S) and XhoI (NEB R3189S) restriction enzymes, and clean-up was performed to remove the non-specific small fragments. Each end of the DNA molecules was biotinylated by ligating with the 500 bp PCR amplified and digested DNA molecules containing many biotins (Biotin-handles). For directionality, Aminoallyl-dUTP-Cy5 (Jena Bioscience NU- 803-CY5-L) was incorporated in one of the 500 bp fragment. The ligation reaction was set-up using T4 DNA Ligase where the DNA end digested with XhoI ligated with a XhoI digested 500 bp biotin-handles, whereas the other end of the molecule digested with NotI was ligated with a NotI digested 500 bp biotin-handle with Biotin-dUTP-Cy5 (Jena Bioscience NU-803-CY5-L) at the start of DNA. The resulted length of the DNA molecule was 21 kb. The ligated product was separated from the non-ligated product by running on the 0.8% Agarose gel. After electrophoresis run, we stained the gel with EtBr and imaged using a fluorescence GelDoc, excised the 21 kb band, and DNA fragment was extracted from it.

### Preparation of 18 kb Cy5-labelled biotinylated DNA

An 18 kb plasmid was prepared by PCR amplification of 9 kb and 8 kb fragment from pCR-XL- 2189-2239 and pCR-XL-2193-2194 plasmid respectively and ligation with 1.5kb Ori and Kanamycin region^42^. The 18 kb plasmid DNA vector was digested with NotI and XhoI restriction enzymes to biotinylated the DNA. Then ligated the digested DNA with already digested 500 bp biotin-handles with the same enzymes using T4 DNA ligase, one was with XhoI-biotin and another one was NotI-cy5 biotin. The ligated product was run on the 0.8% agarose gel to get rid of non-ligated fragments by excised and extracted the desired product only from the gel using Qiagen gel-extraction kit. The end-product was 18 kb DNA with Cy5 signal at one end, similar to the 3 kb (middle) high AT-content 21 kb Cy5-labelled biotinylated DNA.

### Preparation of 21kb DNA with 3 kb moderate AT-content as a control

A 21 kb DNA was prepared in the same way 3 kb (middle) high AT-content 21 kb Cy5-labelled biotinylated DNA was prepared. In this DNA construct, a 3 kb DNA region with 44% (moderate) AT-content was amplified and incorporated PciI and XbaI using 3 kb AT+XbaI_FP forward primer and 3 kb AT+PciI_RP reverse primer from a DNA vector. This 3 kb region has the similar AT-content to its backbone 18 kb DNA vector. The amplified fragment was gel-extracted and digested with PciI and XhoI restriction enzymes. The digested product was cloned into a 18 kb DNA vector. The positive clone was amplified and prepared the biotinylated DNA molecules in similar way we have done for above constructs using XhoI and NotI-Cy5 biotinylated handles.

### Preparation of 3kb AT-rich segment in 21kb DNA at the Cy5-free end

A Cy5-labelled biotinylated 21 kb DNA was prepared by inserting a 3 kb 66% AT-region at end of the 18 kb vector. This is the same 3 kb AT-rich segment in 21 kb Cy5-labelled biotinylated DNA but in this construct the region was inserted at one end of the DNA. The 3 kb AT-rich region was amplified and incorporated KasI + ApaI site at one end and XhoI site at another end using 3+1.5kb end ApaI+KasI FP as forward primer and 3kb+1.5kb_end XhoI_RP as reverse primer respectively. The 18 kb vector was digested with ApaI and XhoI, and 3 kb DNA was also digested with the same restriction enzymes and then cloned it. The positive clone was digested with ApaI and KasI enzyme, then ligated with already digested 500 bp biotin handles with specific restriction sites, one end with ApaI-biotin_cy5 handle and another end with kasI-biotin handle. The ligated product was run on the 0.8% agarose gel and extracted the desired product. The end-product was 21 kb DNA with 3 kb high AT-content region at one end, opposite end of the cy5 signal on the DNA.

### Preparation of 4.8 kb and 1.8 kb DNA

The linear 1.8 kb and 4.8 kb DNA was formed by the digestion of a 6.6 kb pMAL-C2X vector (Addgene 75286) with HindIII (NEB R3104) and EcoRV (NEB R3101) restriction enzymes, giving 1.8 kb and 4.8 kb DNA. The digested reaction mixture was running on the 1% agarose gel, and then DNA bands were seen at 1.8 kb and 4.8 kb. Both DNA bands were excised and gel-extracted using a gel-extraction kit (Qiagen 28706).

### Total Internal Reflection Microscopy

The imaging was performed on an objective-type total internal reflection fluorescence microscope (TIRFM). This microscope is equipped with a Nikon Ti2 eclipse microscope, a motorized H-TIRF, a perfect focus system (PFS), and a Teledyne Photometrics PRIME BSI sCMOS camera. The imaging was done by using an oil immersion objective lens (Nikon Instruments Apo SR TIRF 100×numerical aperture 1.49, oil) under Total Internal Reflection conditions. The sample was illuminated with lasers produced by an L6cc laser combiner from Oxxius Inc., France. For Imaging, 2 × 2 binning of pixels was used with the camera cropped to an effective size of 512 × 512 pixels, where each pixel covered an area of 130 × 130 nm. The acquisition was made by setting the camera to a readout sensitivity of 16 bits.

### Preparation of PEG-passivated Slides

Microscopy slides with microfluidic flow channels were prepared in-house as described previously. For this, holes were drilled on each side of the glass slide (VWR 631-1550) using a Dremel drill (Meisinger 801-009-HP) with a diamond head drill bit of 2 mm diameter for the preparation of microfluidic channels. To reuse the slides, firstly slides were cleaned in a series of steps, initially sonicating the slides in 10% v/v dishwashing detergent for 20 minutes, then rinsing and sonicating in MilliQ water for 5 mins. Placed the slides in Teflon racks and kept them in a beaker sonicating in Acetone (EMPARTA ACS 1.07021.2521) for 10 mins. Then, slides and coverslips (VWR 631-0147) were sonicated for 25 mins in 1 M KOH (Potassium Hydroxide, SRL 84749) solution for etching to generate the hydroxyl groups on the surface, then replaced the KOH with MilliQ water and washed, followed by sonication for 10 mins. Piranha etching was performed on the slides and coverslips for further cleaning. It was prepared by slowly adding 30% H_2_O_2_ (EMPLURA 107209) to H_2_SO_4_ (Fisher Scientific 29997) in a 1:3 volume ratio. Mix the solution properly by stirring the Teflon holders and leave for 30 mins until reaction stops and it stops boiling. The piranha solution was disposed and replaced and washed with water then slides and coverslips were washed with methanol thoroughly and sonicate for 20 mins.

After this Aminosilanation were performed, where mixture was prepared with 1:2:20 ratio of glacial acetic acid (EMPARTA ACS 1.93002.2521), APTES (3-aminopropyl triethoxysilane, TCI A0439), and methanol (EMPARTA ACS 1.07018.2521), respectively. Poured that freshly prepared mixture onto the slides and coverslips then incubated for 25 mins. In this step slides and coverslips were functionalizing with an amine group. After this, washed the slides and coverslips at least 5 times thoroughly with methanol. Each slide or coverslip was p washed thoroughly with MilliQ water and then dried with compressed nitrogen gas.

After amino salinization surface were PEG-passivated using a mixture of biotin-PEG-SVA (Succinimidyl valerate) (Laysan Bio Biotin-PEG-SVA-5000) to mPEG-SVA (Laysan Bio mPEG-SVA-5000) in 1:10 ratio. This ratio of compounds was added in freshly prepared 0.1 M of K_2_SO_4_ (Potassium sulphate) buffer of pH 9-9.5, to reach the final concentration was 20 mM. A 60 µl of this mixture was then sandwiched between a glass slide and a coverslip and incubated overnight in dark and humid environment. Next day, the slides and coverslip were disassembled, rinsed with MilliQ water, and dried using compressed nitrogen gas. An additional round of PEGylation was carried out for further passivation on the surface before using it for single-molecule imaging experiments. For this purpose, we used MS(PEG)_4_ (Thermo Scientific 22341). A 7 µl of 250 mM MS(PEG)_4_ was dissolved in 63 µl of K_2_SO_4_ buffer.

### Single-molecule assay for experiments

Microfluidic flow channels were prepared by sandwiching the double-sided transparent tape between the quartz slide and coverslip. The inlet and outlet holes were made for buffer exchange and for the samples to be applied in the channel. The volume of the channel is approximately 10 µl. The inner surface of the slide and coverslip was PEG-passivated with a 20% (w/v) solution of PEG and biotin-PEG in a 50:1 ratio dissolved in a freshly prepared 100 mM potassium sulphate buffer to block the non-specific bindings and provide a platform for biotin-streptavidin interaction on the surface. For more passivation and blocking of the microfluidic flow channel, first we incubated 10% BSA (Bovin serum albumin, GBiosciences RC1021) for 10 minutes and washed the unbound with 800 µl of T50 buffer (50 mM Tris-HCl pH 7.5, and 50 mM NaCl, 1 mM EDTA), then incubated 0.1% Tween 20 (Sigma P9416) for 5 minutes and washed with 800 µl of T50 buffer. To immobilize the DNA on the surface, we applied 20 µl of 0.1 mg/ml neutravidin (Sigma 31000) in T50 buffer for 10 minutes. Then, to remove the excess unbound streptavidin, it was washed with 800 µl of T50 buffer.

A 30 µl of biotinylated-DNA of approximately 5-10 pM flowed into the flow channel with a constant flow rate of 6-15 µl/min with the help of a motorized syringe pump on the PEG passivated surface to immobilize the DNA through biotin-neutravidin-biotin linkage on the surface. The DNA with biotin either at both ends or at one end binds to the surface, whereas the DNA molecules without biotin at either end or excess unbound were removed from the surface by 100 µl of T50 buffer wash. It gave single and double-tether DNA molecules with different contour lengths, depending on the flow rate. The tether DNA molecules on the surface were visualized using an imaging buffer containing the T50 buffer along with DNA-Intercalating dye, either 50 nM of Sytox Orange (Thermo Scientific S11368) or 50 nM of YOYO-1 (Thermo Scientific Y3601) and an oxygen scavenging system (1× PCD, 1×PCA and 1×trolox). The DNA was illuminated using 561 nm or 480 nm wavelength of lasers to excite Sytox orange or YOYO-1, respectively. The protein Lsr2-Cys labelled with Cy3 or Cy5 was illuminated with 561 nm or 640 nm wavelength of lasers.

40× PCA solution (Protocatechuic acid/ 3,4-Dihydroxybenzoic acid; Sigma 37580-100G-F) was prepared by dissolving 154 mg of PCA in 10 ml of HPLC-grade water adjusted to pH 9.0 using 1 M NaOH. PCD (protocatechuate 3,4 dioxygenase) stock was made by dissolving in a buffer containing 100 mM Tris-HCl pH 8.0, 50 mM KCl (Sigma P9541-1KG), 1 mM EDTA (SRL 35888), and 50% glycerol to make 20× PCD with a final concentration of 6 µM. Trolox (6- hydroxy-2,5,7,8-tetramethyl chroman-2- carboxylic acid; Sigma 238813-1G) was dissolved in 345 µL of 1M NaOH, 430 µL of methanol, and 3.2 mL of HPLC grade water to make 100X Trolox.

### Data Analysis

Data was collected using NIS Elements software and converted to TIFF files. The files were then analyzed using a custom-built Matlab (Mathworks) software. Fluorescence intensity profiles of DNA molecules exhibiting compacted structures were obtained by summing the intensity values from 11 pixels taken across a line perpendicular to the extended DNA in each frame. Background intensity was removed by 2d-median filtering. After normalizing and obtaining the intensities for all frames, the intensity profiles taken at subsequent time points were concatenated to build an intensity kymograph. From the kymographs obtained from individual DNA molecules, the amount of DNA in the compacted regions (in bp) was estimated as follows.

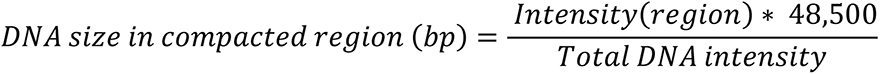

For the analysis of the 21 kb DNA molecules containing an AT-rich region in the middle, individual DNA molecules were initially collected in ImageJ. From these snapshots, a time-averaged projection was generated by averaging the DNA intensity over 200 frames. Additionally, all end-to-end distances of the molecules were normalized between 0 and 1. This normalization enabled us to determine the location of the condensate along the DNA. Subsequently, the data was analysed, interpolated, and averaged to generate the plot depicted in the results section.

### Phase separation assay

All phase separation experiments were carried out by mixing the required concentration of Lsr2 protein with either 4.8 kb or 1.8 kb dsDNA in a buffer containing 50 mM Tris-HCl (pH 7.5) and 50 mM NaCl, 1 mM EDTA and 5 % PEG.

These experiments were performed in two sets: one with different concentrations of protein alone and another set with different concentrations of protein mixed with 0.6 nM of 4.8 kb and 1.8 kb linear DNA. The reaction mixtures were incubated for 15–20 minutes at 4°C and then transferred to a flow cell for imaging. The imaging was carried out using a confocal microscope for the bulk FRAP experiments and a Nikon Eclipse Ti2-E microscope for the quantifying the condensate area and the single-molecule FRAP experiments.

### Analysis of condensate area

To quantify the condensate area under different conditions, multiple fields of view (FOVs) were taken. Background subtraction was applied to the images, and an appropriate thresholding method was set automatically using ImageJ. The binary images were processed, and the particle analysis function was used to segment and measure the condensate areas.

### FRAP analysis

Fluorescence microscopy was performed using Olympus FV 300 Confocal microscope equipped with a 100x oil immersion objective (pixel size: 62 nm). A small region inside the droplets or the entire droplet of Lsr2 condensates and Lsr2-DNA co-condensates labeled with Cy3-Lsr2-Cys or Cy5-Lsr2-Cys proteins was selectively bleached using a 561 nm or 640 nm laser.

Post-bleaching, fluorescence recovery was monitored by capturing images every 1 second. The fluorescence recovery after photobleaching (FRAP) data was analyzed using ImageJ software to generate plots of fluorescence recovery over time.

### FRAP with single-molecule experiments

10 pM of biotinylated λ-DNA was applied to the passivated glass-surface with a flow rate of 6-7 µl/min, visualized the DNA with Sytox Orange fluorescence dye. 500 nM Cy5-labelled Lsr2-Cys protein was applied in the flow channel with a 10 µl/min flow rate. Lsr2 was started to compact the DNA as the DNA was visualized through 561 nm laser and Cy5-Lsr2 in 640 nm laser. The protein was imaged with 1 milliwatt (mW) constant power of 561 nm laser for 100 milliseconds frame rate, including the time gap of 1 second between two consecutive frames. For photobleaching, the power of the laser was increased to 100 mW for 1 sec to bleach the whole field of view, then again reduced the power of the laser and applied the fresh same concentration of the protein for 10 seconds and observed the recovery of the fluorescently labelled Lsr2 protein.

### Analysis of double-tethered DNA experiments

To analyse the DNA sequence inside condensate, we start the analysis by loading microscopy snapshots for DNA molecules. The DNA strand in snapshots is kept parallel to the longitudinal axis and DNA orientation is determined using Cy5 signals. We then subtract the background intensity and generate a line profile by summing pixel intensities perpendicular to the DNA strand. Subsequently, we do the same for all the snapshots for individual molecules and plot a frequency distribution plot (Supplementary Figure 5b & 5c) for the intensities present in the line profiles. From the distribution, we define a threshold as

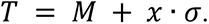

Here, T represents the calculated threshold, M represents the median of the frequency distribution of intensities in intensity profiles, *σ* represents the standard deviation of the distribution, and x is a variable that adjusts the threshold. Here, we set x to 0.5 for proper identification of the condensate region (Supplementary Figure 5d & 5e). Next, we analyze the intensity line profile of individual snapshots to identify the condensate region based on threshold. Indices with intensities higher than the threshold are assigned to condensate regions, while those below the threshold are defined as part of bare DNA. Based on intensities in line profile, we calculate the DNA content for each index on the line profile by

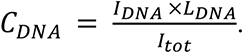

Here, *C_DNA_* is the DNA present in the condensate, *I*_DNA_ represents the intensity of the index, which is detected as part of the condensate, *L_DNA_* is the length of DNA in base pairs and *I_tot_* is the sum of all the intensities in the line profile.

Next, we associate probability of 1 or 0 to base pairs based on their presence or absence in the identified condensate region. This procedure is followed for more than 75 snapshots for each molecule to generate average probability profile. Finally, the probability profiles of many molecules with identical sequence are averaged to generate an average probability profile (Figure 4c & 4g).

### Analysis of DNA envelope width

We estimate the envelope width of single DNA molecules using a method outlined in previous studies^20,24^. As described in the analysis pipeline, first, we generate the time-averaged projections of the DNA signal with and without proteins. After subtracting the background intensity, we extract a line profile from the DNA intensity signals orthogonal to and from middle of the strand. Next, we fit a gaussian curve to the extracted line profile from which the standard deviation is calculated for each molecule. The DNA envelope width is defined as twice the standard deviation calculated from the fitted gaussian curve.

### Simulation methodology

Based on the experimental observations, we employ Brownian dynamics simulations in an NVT ensemble. We integrate the equations of motion at constant temperature of 1.0 *k_B_ T* using a Langevin thermostat. The damping factor (*γ_damp_*) is set to 0.1 𝜏^−1^, and we use a time step of 𝛥𝑡=0.01 𝜏 for integration. We perform these simulations using the ESPResSo package^43^. Further details of the model are discussed in model section.

For simulations with the untethered DNA molecules, we introduced DNA of length 1.8 kb (Figure 1f) and 10 kb (Figure 5a) in a cubic box of dimensions 1.4 μm × 1.4 μm × 1.4 μm. This choice of box dimensions gives us a DNA concentration of 6 *n*M, which recapitulates experiments done with 1.8 kb DNA. For simulations with 1.8 kb, we started simulation with isolated DNA molecules for ∼10^7^𝜏 which is sufficient for the equilibration of the system. Next, we introduced Lsr2 molecules at concentrations of 0.5, 1, 5 and 10 𝜇M as used in the experiments. The DNA-Lsr2 system is then equilibrated for another ∼ 10^7^𝜏. To capture the bulk assay, we simulated the system in absence of the DNA for ∼10^7^𝜏 at Lsr2 concentrations used in the experiments.

To model bridging and co-condensation of Lsr2 and DNA, we introduced a 10 kb long DNA molecule in a cubic box of dimensions mentioned above. Subsequently, we equilibrated the DNA for ∼10^7^𝜏. Next, we introduced Lsr2 at concentration 1 𝜇M and simulate the system for ∼10^7^𝜏.

For simulations with the double-tethered DNA molecules (Figure 4a & 4e), we introduced different 18 kb and 21 kb DNA constructs used in the experiments, in a cuboidal box with dimensions of 1.36 μm × 1.36 μm × 5.1 μm. To minimize the polymer’s interaction with the walls, we placed the DNA molecule at the center of the box and parallel to the longitudinal axis. The distance between the ends of the DNA was set to 3 μm for the 18 kb DNA construct and 3.5 μm for the 21 kb DNA construct by fixing the coordinates of the first and last monomer throughout the simulations. First, we equilibrated the DNA molecule in the absence of Lsr2 for 10^7^𝜏. Following that, we simulated the system for an additional 10^7^𝜏 to generate successive DNA configurations. These DNA configurations were then used as a starting point for different replicate simulations with Lsr2 molecules This procedure ensures unbiased and independent initial conditions for replicate simulations. Next, we introduced Lsr2 molecules in the system at 0.5 𝜇M concentration, and equilibrate for ∼10^7^𝜏.

To generate configurations, we kept the production time to 5 × 10^3^𝜏 for all the simulations mentioned above. By doing this, we ensured equal sampling of the configuration space.

### Model parameters

In the simulations, the interaction strength among Lsr2 dimers was set to 4 *k_B_ T*. This binding energy captured the experimental results observed in Lsr2 dimer phase separation in the absence of DNA (Figure 1f). For modelling experiments with 18 kb and 21 kb DNA constructs, we varied the interaction energy between DNA and Lsr2 from 3 *k_B_ T* to 5 *k_B_ T*, which ensured preferential binding of Lsr2 to AT-rich regions. In the model, AT-content for a 10 bp motif determines the interaction strength between DNA motif and Lsr2. The binding strength is at 3 *k_B_ T* if the AT content is 0%, and increases with a power law to 5 *k_B_ T* as the AT content approaches 100%. Many previous studies with prokaryotes and eukaryotes suggest DNA-protein binding affinity in a range of 0 to 5 *k_B_ T*^21^, which is consistent with our model. Choice of parameters and model details are discussed in the model section.

### Analysis of double-tethered DNA in simulations

For simulations with double-tethered 21 kb DNA constructs, we visualized the simulations using VMD^44^ to count number of condensates present in the system. Next, we initiated the analysis by loading system configurations. We then binned the DNA monomers along the longitudinal axis of the simulation box with a bin size of 3.4 𝑛𝑚, as the DNA strand is parallel to this axis. This gives us a line profile analogous to the intensity line profile generated by summing the intensities that are distributed along the orthogonal rows from DNA snapshots in experiments. For defining the condensate region, we selected a threshold of 2 (as most bins corresponding to bare DNA contain one monomer) and searched for 𝑛 longest stretches in the line profile. Here 𝑛 is defined as number of condensates present in the system. Following that, for each monomer, we associate a probability of 0 and 1 based on their absence or presence in the identified stretches respectively (Supplementary Figure 5g & 5h). For each simulation, we calculated probabilities of monomers from last 500 configurations and averaged them to create probability profiles. Furthermore, for different replicate simulations of the same DNA construct, the probability profiles were averaged and reported (Figure 4d & 4h).

