## Supplementary Information for "Sequence-dependent co-condensation of Lsr2 with DNA elucidates the mechanism of genome compaction in *Mycobacterium tuberculosis*"

**A simple theoretical model to study Lsr2-DNA co-condensation**

We use a minimal model to study Lsr2-DNA co-condensation, as described in an earlier study[1]. We represent the DNA as a coarse-grained semiflexible polymer composed of $n$ monomers, each corresponding to 10 base pairs (bps) in length. Here $n$ depends on the length of the various DNA constructs used in the experiments. We represented Lsr2 dimers as spherical beads with the same size as one monomer along the DNA, since each Lsr2 dimer binds to 10 bps of DNA. Harmonic springs are used to link the adjacent monomers, whereas harmonic angle potential between consecutive bonds captures the semi-flexible nature of DNA. The bond length potential is given by

$$U(r) = \frac{1}{2} k_b (r - r_0)^2 \ .$$

Here, $k_b$ and $r_0$ are the energy corresponding to bond length and equilibrium bond length, respectively. We set $k_b = 100\, k_B T \sigma^{-2}$ and $r_0 = 3.4\ nm$, where $k_B$ is Boltzmann's constant, $T$ is absolute temperature, and $\sigma$ is the unit in length scale (σ = 3.4 nm). The bond angle potential is given by

$$U(\theta) = \frac{1}{2} k_\theta (\theta - \theta_0)^2 \ .$$

Here, $k_\theta$ and $\theta_0$ are the energy corresponding to the bond angle, and equilibrium bond angle, respectively. We set $k_\theta = 15\, k_B T$ and $\theta_0 = 180$, where $k_\theta$ is related to the persistence length ($l_p$) of DNA (150 bps) as

$$l_p = \frac{k_\theta}{k_B T} \ .$$

To model DNA as a self-avoiding polymer, we use Weeks-Chandler-Anderson (WCA) potential between the monomers present in DNA. This potential is a purely repulsive form of Lennard-Jones potential. The interactions between Lsr2 dimers and DNA monomers, as well as between the dimers themselves, are modeled using a Lennard-Jones potential with an attractive tail

$$U_{nb}(r_{ij}) = \begin{cases} 4\varepsilon_{ij} \left[ \left(\frac{\sigma}{r_{ij}}\right)^{12} - \left(\frac{\sigma}{r_{ij}}\right)^{6} + \frac{1}{4} \right], & r_{ij} \le r_{cut} \\ 0. & r_{ij} > r_{cut} \end{cases}$$

Here $r_{ij}$ is the distance among the interacting particles $i$ and $j$, and $r_{cut}$ represents the cutoff distance. It is set to 2.5 $\sigma$ (σ = 3.4 nm) for Lsr2-Lsr2 and DNA-Lsr2 interactions (attractive interaction), whereas it is $2^{1/6}\sigma$ for interactions among the DNA monomers (repulsive interaction). $\varepsilon_{ij}$ represents the interaction strength between particles $i$ and $j$.

We set the interaction strength for Lsr2 dimers within themselves to 4 $k_BT$ for all simulations. The choice of interaction strength between Lsr2 molecules was chosen in a manner to recapitulate the experimental observations obtained from the bulk assay. However, we vary the interaction energies between DNA monomer and Lsr2 to capture various experimental observations, as noted below.

For simulations modeling the experiments done with 1.8 kb DNA molecules, we introduced the DNA as a homogeneous polymer. Here, all the monomers interact with Lsr2 molecules with the interaction strength of 4 $k_BT$ which is the same as interaction strength between protein-protein interactions.

To model DNA-protein interactions in the single-molecule assay, we introduced a power law relationship between AT content of the 10 bp motif and the strength with which it interacts with Lsr2 molecules, given by

$$\varepsilon_{MP} = \varepsilon_0 + 2 \cdot (C_{AT})^{10}.$$

Here, $\varepsilon_{MP}$ represents the interaction strength between DNA monomer and protein, $\varepsilon_0$ is affinity of Lsr2 to the monomers. For 10 bps motifs that does not contain any adenine or thymine, we set the Lsr2-monomer affinity to 3 $k_BT$. $C_{AT}$ represents the fraction of AT content in 10 bp-long motifs. Depending on the AT content, interaction strength varies from 3 $k_BT$ to 5 $k_BT$ (Supplementary Figure 6b). We summarize the monomer-protein interaction parameters in Table 1. We keep protein-protein interactions to 4 $k_BT$ as in the previous sequences.

**Table 1**. AT content of 10 bps of eleven kinds of monomers in the double-tethered DNA models, and their interaction affinities with Lsr2 dimers.

| $AT_{10}$ | $\varepsilon_{MP}(k_BT)$ |
|---|---|
| 0.0 | 3.00 |
| 0.1 | 3.00 |

| | |
|---|---|
| 0.2 | 3.00 |
| 0.3 | 3.00 |
| 0.4 | 3.00 |
| 0.5 | 3.00 |
| 0.6 | 3.01 |
| 0.7 | 3.06 |
| 0.8 | 3.21 |
| 0.9 | 3.70 |
| 1.0 | 4.99 |

Finally, to model a 10 kb DNA, as described in the Discussion section, we represented the DNA as a block copolymer consisting of two types of monomers A and B. Here, five 30-monomer blocks of monomer type A are separated by four 170-monomer blocks of monomer type B. Monomer type A binds Lsr2 with interaction strength of 5 $k_B T$ whereas for monomer type B, the interaction strength is 0.1 $k_B T$.

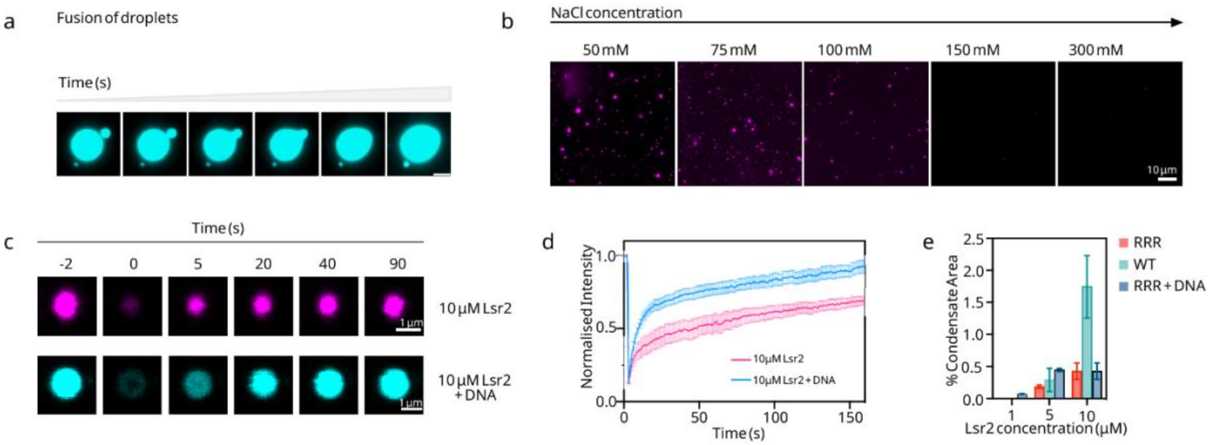


**Supplementary Figure 1: Lsr2 and DNA undergo liquid-liquid phase-separation.** a) Fusion event of two phase separated droplets (scale bar – 3 µm). b) Lsr2-DNA co-condensates dissolve when concentration of salt is increased (3 µM Lsr2 + 2ng of 4.6 kb DNA) (scale bar – 10 µm). c) Fluorescence images showing the photobleaching of 10 µM Lsr2 droplets (above) 10 µM Lsr2 in the presence of DNA (scale bar – 1 µm). d) Plot showing the normalized fluorescence intensity recovery after bleaching an entire droplet. e) Condensate area quantification of WT and RRR mutant.

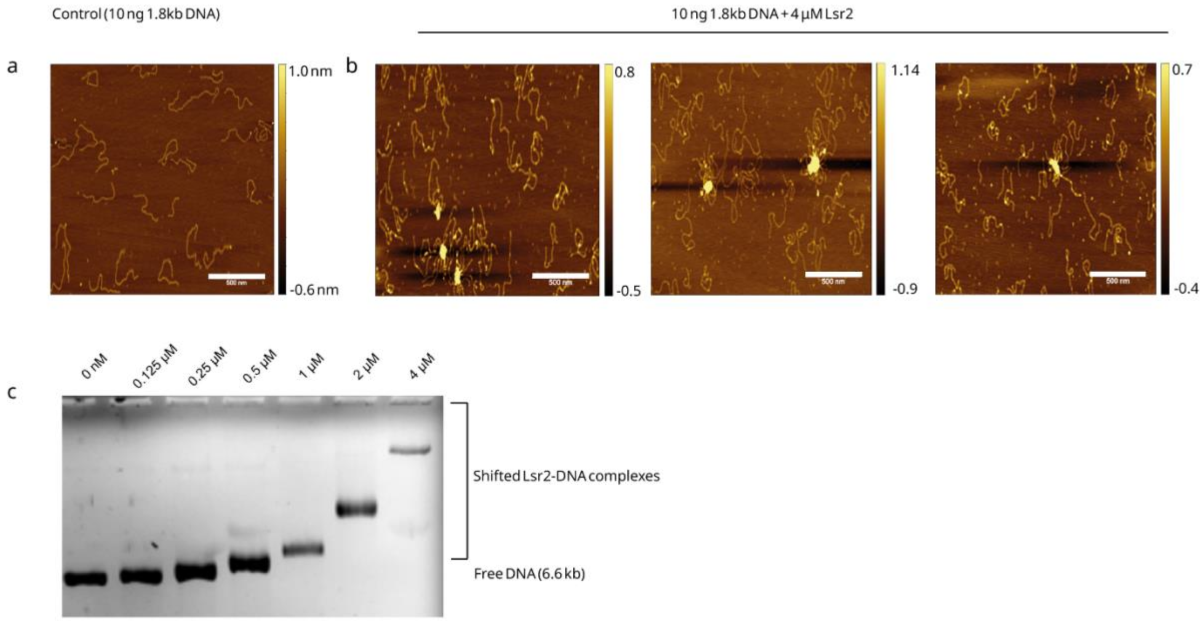


**Supplementary Figure 2: Lsr2 bridges DNA and forms nucleoprotein complexes.** a) Atomic Force Microscopy (AFM) images of 10 ng of 1.8 kb linear double stranded DNA and b) DNA-Lsr2 complex at 4 µM Lsr2 concentration. c) 1% agarose gel image after performing EMSA with 6.6 kb DNA. Scale bar – 500 nm (a and b)

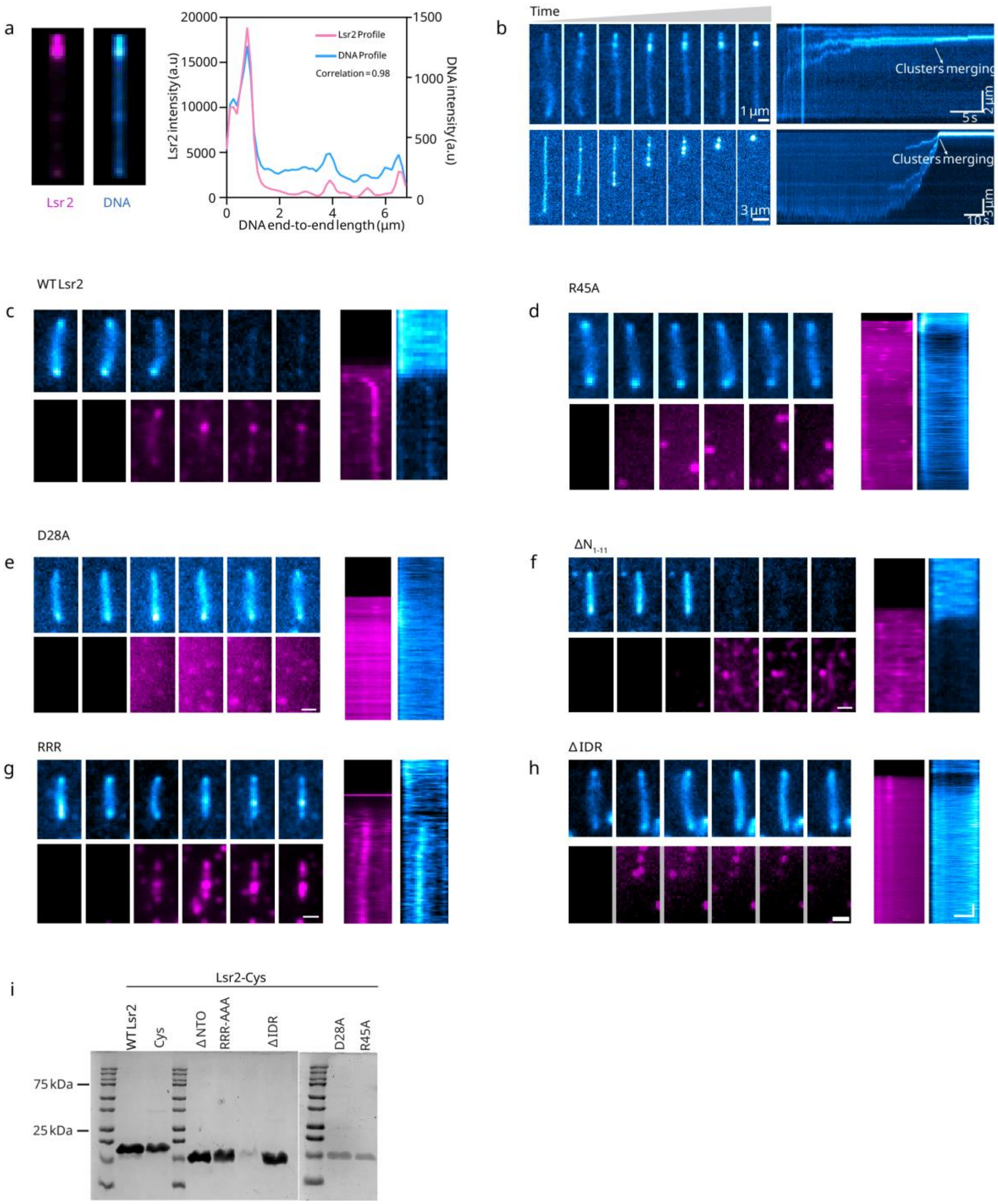


**Supplementary Figure 3: Single-molecule imaging of Lsr2 mutants.** a) Correlation analysis reveals a co-localization between the intensity peaks of Lsr2 and those of the DNA molecule, indicating that Lsr2 and DNA form clusters together b) Snapshots depicting the merging of clusters over time on a double tethered DNA (top) and a single tethered DNA (bottom). c) Series of snapshots representing the single-molecule experiments (left) and the corresponding kymograph (right) for WT Lsr2, d) R45A, e) D28A, f) ΔNTD, g) RRR, and h) ΔIDR. (DNA is represented in blue and protein variant in magenta) i) SDS-PAGE gel image of purified WT Lsr2 along with all the mutants.

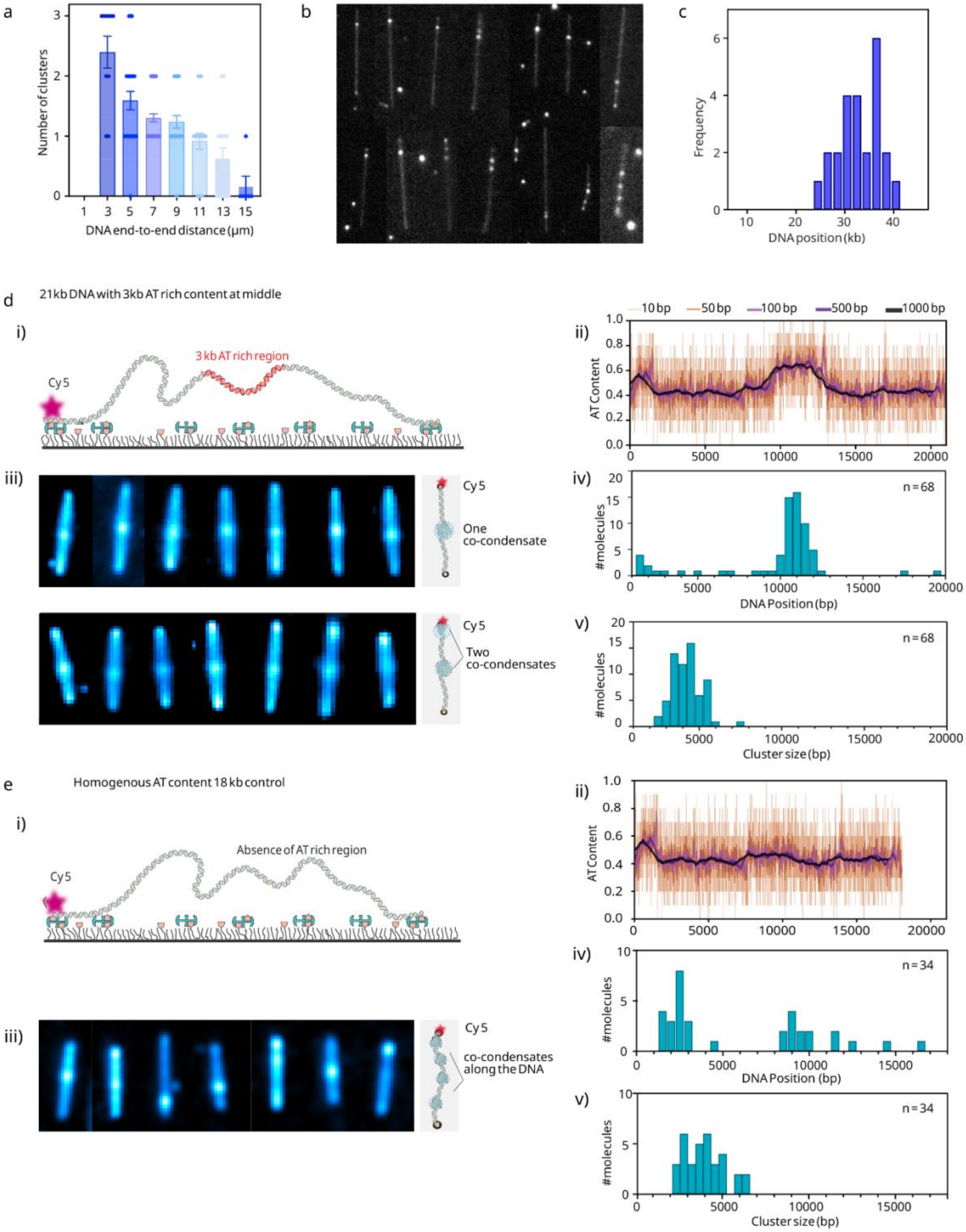

a
Number of clusters
DNA end-to-end distance (μm)
b
c
Frequency
DNA position (kb)
d
21kb DNA with 3kb AT rich content at middle
i)
Cy5
3 kb AT rich region
ii)
10 bp
50 bp
100 bp
500 bp
1000 bp
AT Content
iii)
One co-condensate
Two co-condensates
iv)
n = 68
#molecules
DNA Position (bp)
v)
Cluster size (bp)
e
Homogenous AT content 18 kb control
Absence of AT rich region
n = 34
co-condensates along the DNA

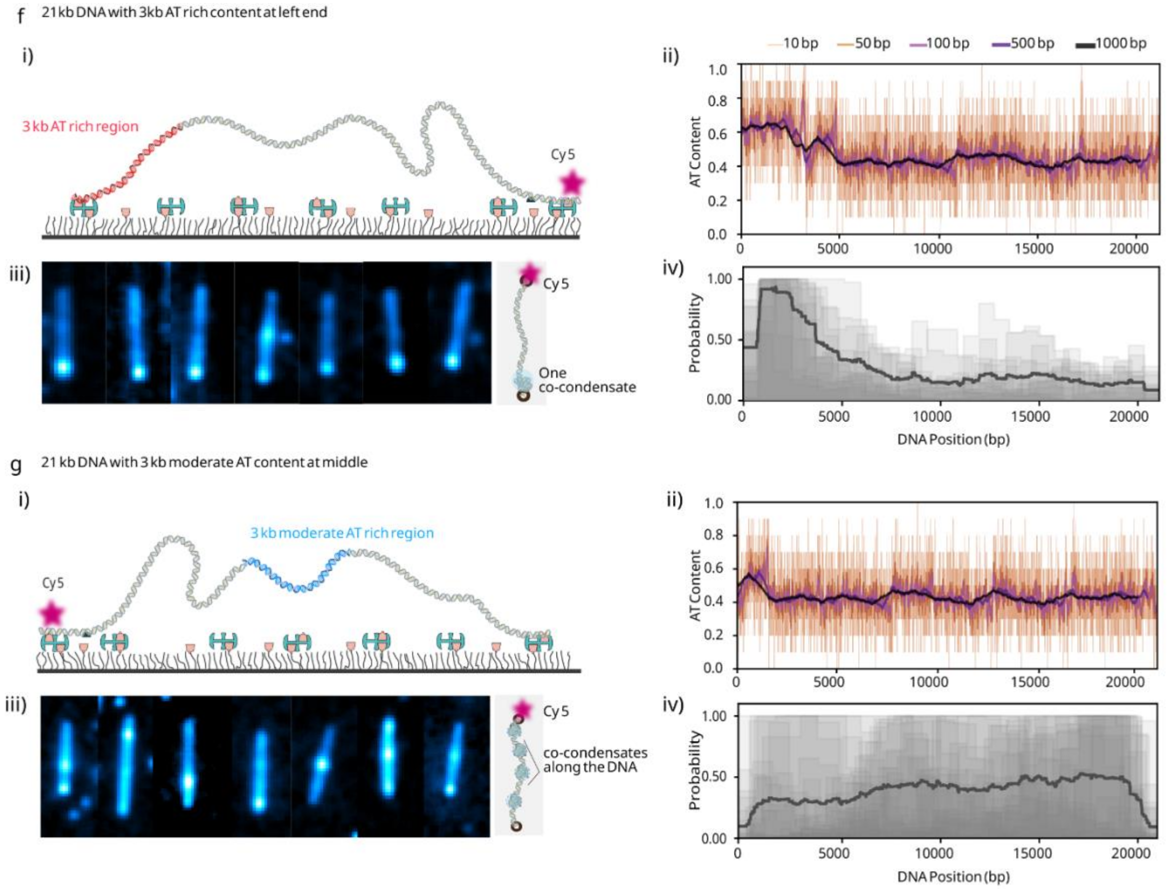


**Supplementary Figure 4: Lsr2-DNA clusters form at specific regions on DNA.** a) Plot depicting the variation of the number of clusters as a function of the DNA end-to-end distance. b) A collage of snapshots showing the clusters forming towards on end of the λ-DNA molecules. c) Histogram showing that the frequency of clusters observed is high towards one end of the λ DNA. d)i) Schematic of Cy5 labelled DNA construct with an AT rich region in the middle. ii) AT profile of the DNA construct in (i) calculated by averaging every 10, 50, 100, 500 and 1000 base pairs. iii) Representative snapshots of various DNA molecules with a co-condensate at the AT-rich center (above) and co-condensates at both the center and the end (below). Two clusters were seen in double tethered DNA with a shorter end to end distance of less than 2.5 μm. iv) Histogram showing that most of the co-condensates (86.4 %) form at the AT rich center (above). Histogram displaying the sizes of the co-condensates found on the DNA molecules (below). e) i) Schematic of Cy5 labelled DNA construct without an AT rich region. ii) AT profile of the construct. iii) Representative snapshots displaying co-condensates at various positions along the DNA. iv) Histogram showing that the co-condensates form at different positions along the length of the DNA(above). Histogram displaying the sizes of the co-condensates found on the DNA molecules (below) f) i) Schematic of the Cy5 labelled DNA with an AT rich region towards the end. ii) AT profile of the DNA construct. iii) Representative snapshots of various DNA molecules with a co-condensate at the AT rich end. iv) Probability profile of the position of cluster formation on the DNA. g) i) -iv) similar as in f) but for a Cy5 labelled DNA construct with moderate AT region in the middle.

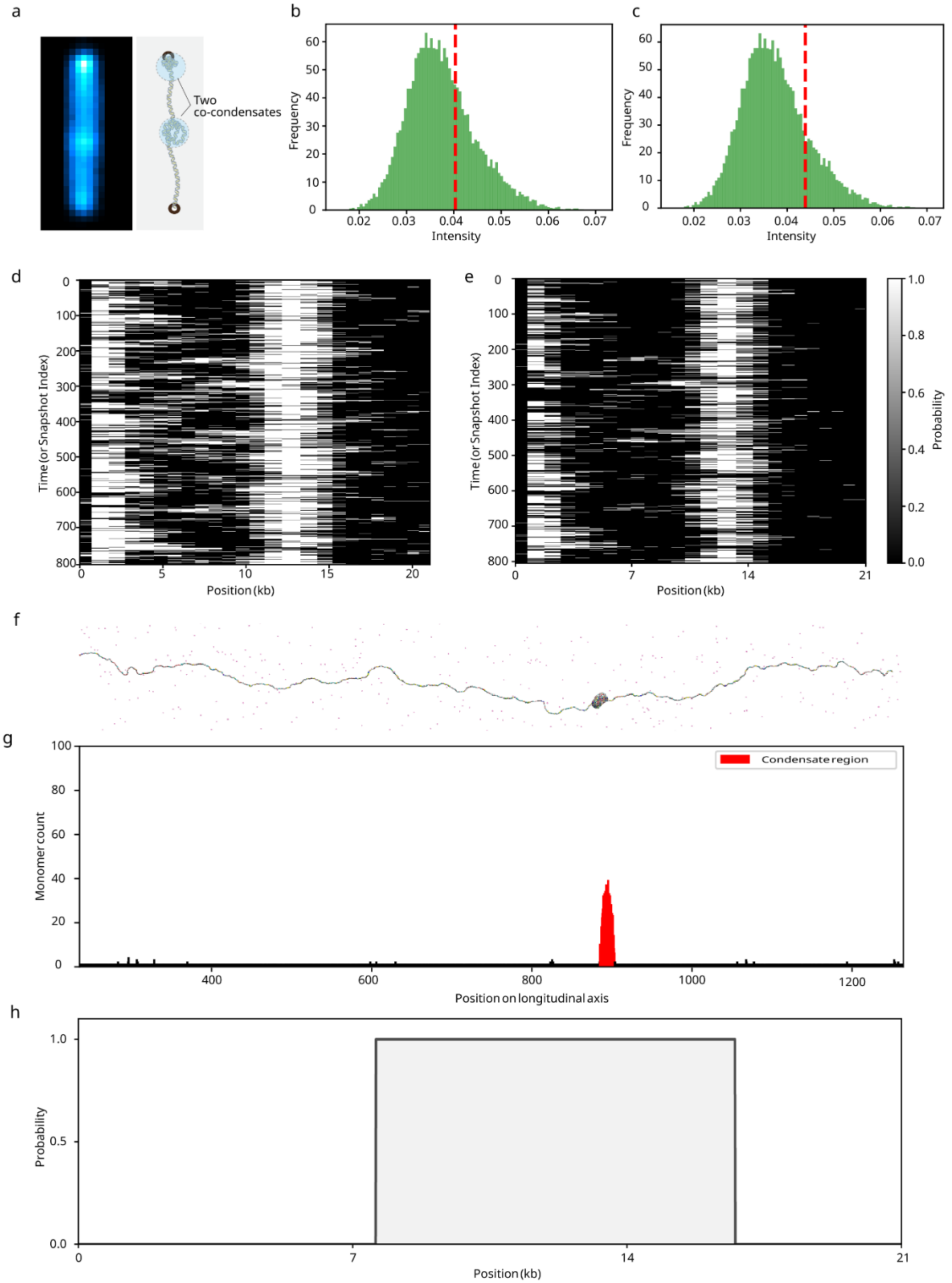
a
Two co-condensates
b
Frequency
Intensity
c
Frequency
Intensity
d
Time (or Snapshot Index)
Position (kb)
e
Time (or Snapshot Index)
Position (kb)
Probability
f
g
Condensate region
Monomer count
Position on longitudinal axis
h
Probability
Position (kb)

**Supplementary Figure 5: Estimation of DNA within the condensates in experiments and simulations.** (a). A time-averaged projection of 21 kb DNA with high AT content in the centre is shown along with a cartoon representation of the molecule. (b) and (c) represent the frequency distribution of normalized DNA intensities present in the line profile generated for the DNA molecule. The red dashed lines represent different thresholds; x, which is a parameter in threshold estimation, is set to 0.5 and 1 respectively (see the Methods section for threshold details). (d) and (e) represent the kymographs corresponding to the two thresholds mentioned above (in order, x = 0.5 and x = 1). f) represents a configuration taken from a simulation with the 21 kb DNA construct with high AT-content in the middle. (g) A histogram is plotted for the monomer count along the axis parallel to the DNA strand. Red bins indicate the DNA in condensate. (h) A probability profile is generated for the DNA based on presence or absence of the monomer in the condensate region.

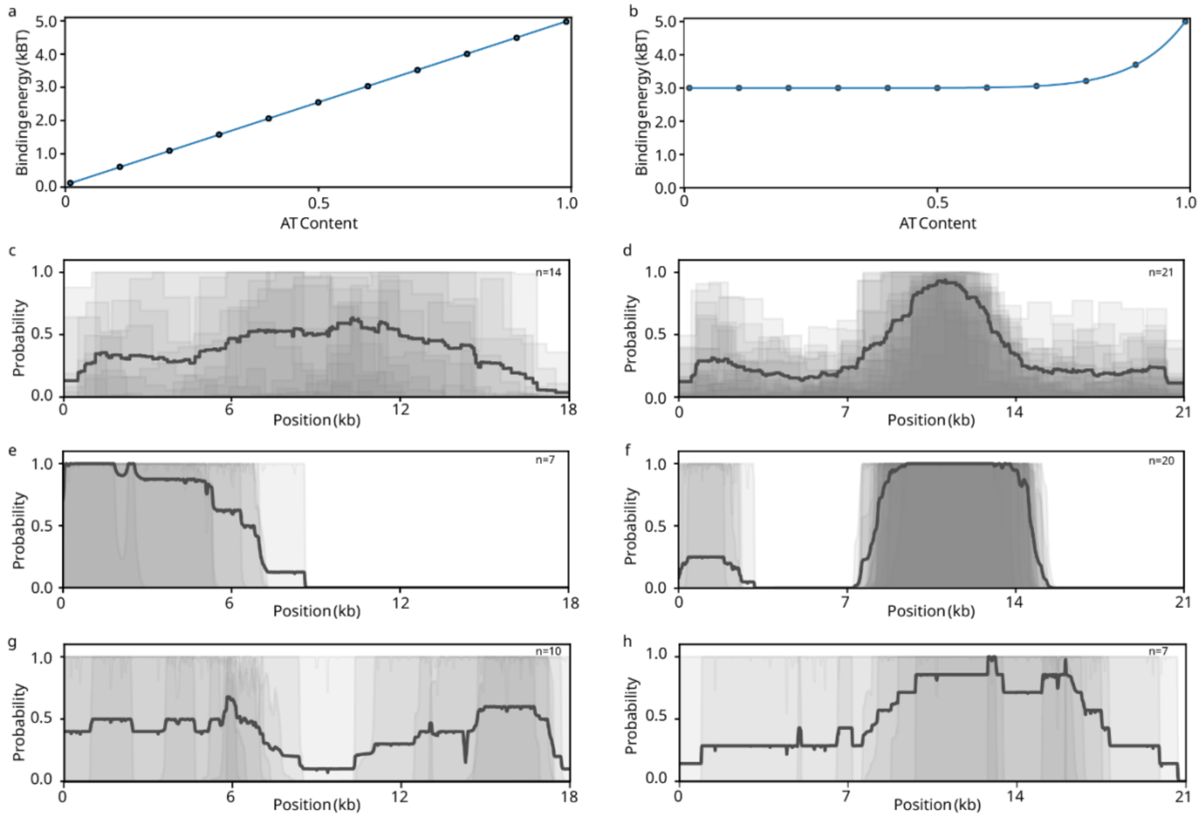


**Supplementary Figure 6: Relationship between binding energy and AT-content (10 bp) of DNA motif in Lsr2-DNA interactions.** (a) A linear relation between the binding energy of Lsr2-DNA interaction and AT-content of the monomer is shown, here the binding energy varies from 0.1 to 5 $k_B T$ depending on the AT-content of the motif (monomer). (b) A power law relation between the binding energy of Lsr2-DNA interaction and the AT-content of the monomer is depicted (see the Model section for details). (c) and (d) The probability profiles of 18 kb DNA (control) and 21 kb DNA with high AT content in the middle, obtained from single-molecule experiments (left to right). The same data is shown in Figure 4. (e) & (f) Probability profiles generated from simulations that follow a linear relation between the binding energy of Lsr2-DNA interaction and AT-content of the monomer (18 kb & 21 kb; left to right, respectively). (g) and (h) represent the probability profiles generated from simulations using a power law relation between the binding energy of proteins to the DNA monomers and AT-content of the respective monomers (18 kb & 21 kb; left to right respectively).

**Table 2: Primers list**

| Sr. no | Construct Name | Primer Name | sequence |
|---|---|---|---|
| 1 | Lsr2_Cys | Lsr2_cys_NcoI_FP | ATTCGACCATGGTATGCGCGAAGAAAGTAACCG |
| | | Lsr2_cys_XhoI_RP | TGGTGGTGCTCGAGGGTCGCCG |
| 2 | Lsr2_ΔNTO | Lsr2_ΔNTO_FP | GGTATATCTCCTTCTTAAAGTTAAA |
| | | Lsr2_ΔNTO_RP | GATTTCGACGGTTCGGGCGC |
| 3 | Lsr2_ΔIDR | Lsr2 trunc. 58-73_FP | GCGACGGCCCGCCGCCACCC |
| | | Lsr2 trunc. 58-73_RP | GCGATCGACCGCGAGCAGAGC |
| 4 | Lsr2_D28A | Lsr2_D28A_FP | GGCAAGCCCGAATTCGACCG |
| | | Lsr2_D28A_RP | GGGGTGACCTATGAGATCGAC |
| 5 | Lsr2_R45A | Lsr2_R45A_FP | CAGTTTCGTGGCATTCTTAG |
| | | Lsr2_R45A_RP | GCCGGCGACCTGAAGCAATG |
| 6 | Lsr2_RRR_AAA | Lsr2_RRR_FP | CCCACCGACGCGACGGCCCG |
| | | Lsr2_RRR_RP | GCGGCCGCGGGCCGTTCCGGATCCGGCCG |
| 7 | 21kb_3kb AT-rich | 3 kb 66% AT+XbaI_FP | TTTTTTTCTAGATCCCTCTGAAAAAATCTTCCG |
| | | 3 kb 66% AT+PciI_RP | TTTTTTTTTTACATGTTGATTTTATTTTGACTGATAGTG |
| 8 | 21kb_3kb_moderate | 3 kb AT+XbaI_FP | TTTTTTTTTTTCTAGAGCAGAGCTGGAAGTGCAGAC |
| | | 3 kb AT+PciI_RP | TTTTTTTTTTACATGTAAACATCCCTTACACTGGTG |
| 9 | 21kb_3kb_end | 3kb_ApaI+KasI_FP | TTTTTTTTTTGGGCCCTCCCTCTGAAAAAATCTTCCGGGCGCCAGTTTGCTAGGCACTGATAC |
| | | 3kb_XhoI_RP | TTTTTTTTTTCTCGAGTGATTTTATTTTGACTGATAG |
| 10 | 500 bp Biotin Handles | BiotinH_FP | GGTTTGCGTATTGGGCGCTCTTCCG |

| | | | |
|---|---|---|---|
| | | BiotinH_RP _KasI | TTTTTTTTTTGGCGCCAGACGATAGTTACCGGATAAGGCG C |
| | | BiotinH_RP _ApaI | TTTTTTTTTTGGGCCCAGACGATAGTTACCGGATAAGGCG C |
| | | BiotinH_RP _NotI | GCGGCCGCAGACGATAGTTACCGGATAAGGCGC |
| | | BiotinH_RP _XhoI | CTCGAGAGACGATAGTTACCGGATAAGGCGC |

**3 kb_ high AT-rich content_21kb DNA Sequence**

TCCCTCTGAAAAAATCTTCCGGGCGCCAGTTTGCTAGGCACTGATACATAACTCTTTTCCAATA
ATTGGGGAAGTCATTCAAATCTATAATAGGTTTCAGATTTGCTTCAATAAATTCTGACTGTAGCT
GCTGAAACGTTGCGGTTGAACTATATTTCCTTATAACTTTTACGAAAGAGTTTCTTTGAGTAAT
CACTTCACTCAAGTGCTTCCCTGCCTCCAAACGATACCTGTTAGCAATATTTAATAGCTTGAAA
TGATGAAGAGCTCTGTGTTTGTCTTCCTGCCTCCAGTTCGCCGGGCATTCAACATAAAAACTG
ATAGCACCCGGAGTTCCGGAAACGAAATTTGCATATACCCATTGCTCACGAAAAAAAATGTCC
TTGTCGATATAGGGATGAATCGCTTGGTGTACCTCATCTACTGCGAAAACTTGACCTTTCTCTC
CCATATTGCAGTCGCGGCACGATGGAACTAAATTAATAGGCATCACCGAAAATTCAGGATAATG
TGCAATAGGAAGAAAATGATCTATATTTTTTGTCTGTCCTATATCACCACAAAATGGACATTTTT
CACCTGATGAAACAAGCATGTCATCGTAATATGTTCTAGCGGGTTTGTTTTTATCTCGGAGATT
ATTTTCATAAAGCTTTTCTAATTTAACCTTTGTCAGGTTACCAACTACTAAGGTTGTAGGCTCA
AGAGGGTGTGTCCTGTCGTAGGTAAATAACTGACCTGTCGAGCTTAATATTCTATATTGTTGTT
CTTTCTGCAAAAAAGTGGGGAAGTGAGTAATGAAATTATTTCTAACATTTATCTGCATCATACC
TTCCGAGCATTTATTAAGCATTTCGCTATAAGTTCTCGCTGGAAGAGGTAGTTTTTTCATTGTAC
TTTACCTTCATCTCTGTTCATTATCATCGCTTTTAAAACGGTTCGACCTTCTAATCCTATCTGACC
ATTATAATTTTTTAGAATGGTTTCATAAGAAAGCTCTGAATCAACGGACTGCGATAATAAGTGG
TGGTATCCAGAATTTGTCACTTCAAGTAAAAACACCTCACGAGTTAAAACACCTAAGTTCTCA
CCGAATGTCTCAATATCCGGACGGATAATATTTATTGCTTCTCTTGACCGTAGGACTTTCCACAT
GCAGGATTTTGGAACCTCTTGCAGTACTACTGGGGAATGAGTTGCAATTATTGCTACACCATTG
CGTGCATCGAGTAAGTCGCTTAATGTTCGTAAAAAAGCAGAGAGCAAAGGTGGATGCAGATG
AACCTCTGGTTCATCGAATAAAACTAATGACTTTTCGCCAACGACATCTACTAATCTTGTGATA
GTAAATAAAACAATTGCATGTCCAGAGCTCATTCGAAGCAGATATTTCTGGATATTGTCATAAA
ACAATTTAGTGAATTTATCATCGTCCACTTGAATCTGTGGTTCATTACGTCTTAACTCTTCATATT
TAGAAATGAGGCTGATGAGTTCCATATTTGAAAAGTTTTCATCACTACTTAGTTTTTTGATAGCT
TCAAGCCAGAGTTGTCTTTTTCTATCTACTCTCATACAACCAATAAATGCTGAAATGAATTCTA
AGCGGAGATCGCCTAGTGATTTTAAACTATTGCTGGCAGCATTCTTGAGTCCAATATAAAAGTA
TTGTGTACCTTTTGCTGGGTCAGGTTGTTCTTTAGGAGGAGTAAAAGGATCAAATGCACTAAA
CGAAACTGAAACAAGCGATCGAAAATATCCCTTTGGGATTCTTGACTCGATAAGTCTATTATTT
TCAGAGAAAAAATATTCATTGTTTTCTGGGTTGGTGATTGCACCAATCATTCCATTCAAAATTG
TTGTTTTACCACACCCATTCCGCCCGATAAAAGCATGAATGTTCGTGCTGGGCATAGAATTAAC
CGTCACCTCAAAAGGTATAGTTAAATCACTGAATCCGGGAGCACTTTTTCTATTAAATGAAAA

GTGGAAATCTGACAATTCTGGCAAACCATTTAACACACGTGCGAACTGTCCATGAATTTCTGA
AAGAGTTACCCCTCTAAGTAATGAGGTGTTAAGGACGCTTTCATTTTCAATGTCGGCTAATCGA
TTTGGCCATACTACTAAATCCTGAATAGCTTTAAGAAGGTTATGTTTAAAACCATCGCTTAATTT
GCTGAGATTAACATAGTAGTCAATGCTTTCACCTAAGGAAAAAAACATTTCAGGGAGTTGACT
GAATTTTTTATCTATTAATGAATAAGTGCTTACTTCTTCTTTTTGACCTACAAAACCAATTTTAA
CATTTCCGATATCGCATTTTTCACCATGCTCATCAAAGACAGTAAGATAAAACATTGTAACAAA
GGAATAGTCATTCCAACCATCTGCTCGTAGGAATGCCTTATTTTTTTCTACTGCAGGAATATACC
CGCCTCTTTCAATAACACTAAACTCCAACATATAGTAACCCTTAATTTTATTAAAATAACCGCAA
TTTATTTGGCGGCAACACAGGATCTCTCTTTTAAGTTACTCTCTATTACATACGTTTTCCATCTA
AAAATTAGTAGTATTGAACTTAACGGGGCATCGTATTGTAGTTTTCCATATTTAGCTTTCTGCTT
CCTTTTGGATAACCCACTGTTATTCATGTTGCATGGTGCACTGTTTATACCAACGATATAGTCTA
TTAATGCATATATAGTATCGCCGAACGATTAGCTCTTCAGGCTTCTGAAGAAGCGTTTCAAGTA
CTAATAAGCCGATAGATAGCCACGGACTTCGTAGCCATTTTTCATAAGTGTTAACTTCCGCTCC
TCGCTCATAACAGACATTCACTACAGTTATGGCGGAAAGGTATGCATGCTGGGTGTGGGGAAG
TCGTGAAAGAAAAGAAGTCAGCTGCGTCGTTTGACATCACTGCTATCTTCTTACTGGTTATGC
AGGTCGTAGTGGGTGGCACACAAAGCTTTGCACTGGATTGCGAGGCTTTGTGCTTCTCTGGA
GTGCGACAGGTTTGATGACAAAAAATTAGCGCAAGAAGACAAAAATCACCTTGCGCTAATGC
TCTGTTACAGGTCACTAATACCATCTAAGTAGTTGATTCATAGTGACTGCATATGTTGTGTTTTA
CAGTATTATGTAGTCTGTTTTTTATGCAAAATCTAATTTAATATATTGATATTTATATCATTTTACG
TTTCTCGTTCAGCTTTTTTATACTAAGTTGGCATTATAAAAAAGCATTGCTTATCAATTTGTTGC
AACGAACAGGTCACTATCAGTCAAAATAAAATCA

**3 kb_moderate_AT-content_21kb DNA Sequence**

GATgAgACGGCCTCTAGAGCAGAGCTGGAAGTGCAGACCGGCATGACACAGCGACGCAGGGG
ACCTGCAGGATTTTATGTATGAAAACGCCCACCATTCCCACCCTTCTGGGGCCGGACGGCATG
ACATCGCTGCGCGAATATGCCGGTTATCACGGCGGTGGCAGCGGATTTGGAGGGCAGTTGCG
GTCGTGGAACCCACCGAGTGAAAGTGTGGATGCAGCCCTGTTGCCCAACTTTACCCGTGGCA
ATGCCCGCGCAGACGATCTGGTACGCAATAACGGCTATGCCGCCAACGCCATCCAGCTGCATC
AGGATCATATCGTCGGGTCTTTTTTCCGGCTCAGTCATCGCCCAAGCTGGCGCTATCTGGGCAT
CGGGGAGGAAGAAGCCCGTGCCTTTTCCCGCGAGGTTGAAGCGGCATGGAAAGAGTTTGCC
GAGGATGACTGCTGCTGCATTGACGTTGAGCGAAAACGCACGTTTACCATGATGATTCGGGAA
GGTGTGGCCATGCACGCCTTTAACGGTGAACTGTTCGTTCAGGCCACCTGGGATACCAGTTCG
TCGCGGCTTTTCCGGACACAGTTCCGGATGGTCAGCCCGAAGCGCATCAGCAACCCGAACAA
TACCGGCGACAGCCGGAACTGCCGTGCCGGTGTGCAGATTAATGACAGCGGTGCGGCGCTGG
GATATTACGTCAGCGAGGACGGGTATCCTGGCTGGATGCCGCAGAAATGGACATGGATACCCC
GTGAGTTACCCGGCGGGCGCGCCTCGTTCATTCACGTTTTTGAACCCGTGGAGGACGGGCAG
ACTCGCGGTGCAAATGTGTTTTACAGCGTGATGGAGCAGATGAAGATGCTCGACACGCTGCA
GAACACGCAGCTGCAGAGCGCCATTGTGAAGGCGATGTATGCCGCCACCATTGAGAGTGAGC
TGGATACGCAGTCAGCGATGGATTTTATTCTGGGCGCGAACAGTCAGGAGCAGCGGGAAAGG
CTGACCGGCTGGATTGGTGAAATTGCCGCGTATTACGCCGCAGCGCCGGTCCGGCTGGGAGG
CGCAAAAGTACCGCACCTGATGCCGGGTGACTCACTGAACCTGCAGACGGCTCAGGATACGG
ATAACGGCTACTCCGTGTTTGAGCAGTCACTGCTGCGGTATATCGCTGCCGGGCTGGGTGTCT
CGTATGAGCAGCTTTCCCGGAATTACGCCCAGATGAGCTACTCCACGGCACGGGCCAGTGCG

AACGAGTCGTGGGCGTACTTTATGGGGCGGCGAAAATTCGTCGCATCCCGTCAGGCGAGCCA
GATGTTTCTGTGCTGGCTGGAAGAGGCCATCGTTCGCCGCGTGGTGACGTTACCTTCAAAAGC
GCGCTTCAGTTTTCAGGAAGCCCGCAGTGCCTGGGGGAACTGCGACTGGATAGGCTCCGGTC
GTATGGCCATCGATGGTCTGAAAGAAGTTCAGGAAGCGGTGATGCTGATAGAAGCCGGACTG
AGTACCTACGAGAAAGAGTGCGCAAAACGCGGTGACGACTATCAGGAAATTTTTGCCCAGCA
GGTCCGTGAAACGATGGAGCGCCGTGCAGCCGGTCTTAAACCGCCCGCCTGGGCGGCTGCAG
CATTTGAATCCGGGCTGCGACAATCAACAGAGGAGGAGAAGAGTGACAGCAGAGCTGCGTA
ATCTCCCGCATATTGCCAGCATGGCCTTTAATGAGCCGCTGATGCTTGAACCCGCCTATGCGCG
GGTTTTCTTTTGTGCGCTTGCAGGCCAGCTTGGGATCAGCAGCCTGACGGATGCGGTGTCCGG
CGACAGCCTGACTGCCCAGGAGGCACTCGCGACGCTGGCATTATCCGGTGATGATGACGGAC
CACGACAGGCCCGCAGTTATCAGGTCATGAACGGCATCGCCGTGCTGCCGGTGTCCGGCACG
CTGGTCAGCCGGACGCGGGCGCTGCAGCCGTACTCGGGGATGACCGGTTACAACGGCATTAT
CGCCCGTCTGCAACAGGCTGCCAGCGATCCGATGGTGGACGGCATTCTGCTCGATATGGACAC
GCCCGGCGGGATGGTGGCGGGGGCATTTGACTGCGCTGACATCATCGCCCGTGTGCGTGACA
TAAAACCGGTATGGGCGCTTGCCAACGACATGAACTGCAGTGCAGGTCAGTTGCTTGCCAGT
GCCGCCTCCCGGCGTCTGGTCACGCAGACCGCCCGGACAGGCTCCATCGGCGTCATGATGGC
TCACAGTAATTACGGTGCTGCGCTGGAGAAACAGGGTGTGGAAATCACGCTGATTTACAGCG
GCAGCCATAAGGTGGATGGCAACCCCTACAGCCATCTTCCGGATGACGTCCGGGAGACACTG
CAGTCCCGGATGGACGCAACCCGCCAGATGTTTGCGCAGAAGGTGTCGGCATATACCGGCCT
GTCCGTGCAGGTTGTGCTGGATACCGAGGCTGCAGTGTACAGCGGTCAGGAGGCCATTGATG
CCGGACTGGCTGATGAACTTGTTAACAGCACCGATGCGATCACCGTCATGCGTGATGCACTGG
ATGCACGTAAATCCCGTCTCTCAGGAGGGCGAATGACCAAAGAGACTCAATCAACAACTGTT
TCAGCCACTGCTTCGCAGGCTGACGTTACTGACGTGGTGCCAGCGACGGAGGGCGAGAACG
CCAGCGCGGCGCAGCCGGACGTGAACGCGCAGATCACCGCAGCGGTTGCGGCAGAAAACAG
CCGCATTATGGGGATCCTCAACTGTGAGGAGGCTCACGGACGCGAAGAACAGGCACGCGTGC
TGGCAGAAACCCCCGGTATGACCGTGAAAACGGCCCGCCGCATTCTGGCCGCAGCACCACAG
AGTGCACAGGCGCGCAGTGACACTGCGCTGGATCGTCTGATGCAGGGGGCACCGGCACCGCT
GGCTGCAGGTAACCCGGCATCTGATGCCGTTAACGATTTGCTGAACAC

**500 bp Biotin Handle**

GGTTTGCGTATTGGGCGCTCTTCCGCTTCCTCGCTCACTGACTCGCTGCGCTCGGTCGTTCGG
CTGCGGCGAGCGGTATCAGCTCACTCAAAGGCGGTAATACGGTTATCCACAGAATCAGGGGAT
AACGCAGGAAAGAACATGTGAGCAAAAGGCCAGCAAAAGGCCAGGAACCGTAAAAAGGCC
GCGTTGCTGGCGTTTTTCCATAGGCTCCGCCCCCCTGACGAGCATCACAAAAATCGACGCTCA
AGTCAGAGGTGGCGAAACCCGACAGGACTATAAAGATACCAGGCGTTTCCCCCTGGAAGCTC
CCTCGTGCGCTCTCCTGTTCCGACCCTGCCGCTTACCGGATACCTGTCCGCCTTTCTCCCTTCG
GGAAGCGTGGCGCTTTCTCATAGCTCACGCTGTAGGTATCTCAGTTCGGTGTAGGTCGTTCGC
TCCAAGCTGGGCTGTGTGCACGAACCCCCCGTTCAGCCCGACCGCTGCGCCTTATCCGGTAAC
TATCGTCT

1 Singh, R. K., Swain, P., Ganji, M. & Choubey, S. Decoding the role of DNA sequence on protein-DNA co-condensation. *bioRxiv*, 2024.2002.2024.581870, doi:10.1101/2024.02.24.581870 (2024).
